# A blastocyst-derived *in vitro* model of the *human* chorion

**DOI:** 10.1101/2025.08.06.668884

**Authors:** Luca C. Schwarz, Matthew J. Shannon, Gina McNeill, Rina C. Sakata, Viviane S. Rosa, Katherine Cheah, Laura Keller, Phil Snell, Leila Christie, Kay Elder, Anastasia Mania, Lauren Weavers, Rachel Gibbons, Tugce Pehlivan Budak, Ippokratis Sarris, Amy Barrie, Alison Campbell, Roser Vento-Tormo, Gary D. Smith, Alexander G. Beristain, Marta N. Shahbazi

**Author notes:** equal contribution (Matthew J. Shannon, Gina McNeill, Rina C. Sakata).

## Abstract

The placenta supports the foetus by mediating nutrient and gas exchange, hormone production, and immune protection. Yet, human placental biology remains poorly understood due to limited *in vitro* models. The foetal part of the placenta, the chorion, consists of a trophoblast-derived layer and a mesenchymal zone containing extraembryonic mesoderm cells. Here, we show that human blastocysts can give rise to both trophoblast organoids and extraembryonic mesoderm cells under the same culture conditions. Trophoblast organoids originate from both outer trophoblast cells and inner cells of the blastocyst, while extraembryonic mesoderm cells derive exclusively from inner cells. These organoids recapitulate the cellular composition, morphology, and function of the *in vivo* trophoblast. Moreover, given the high rate of aneuploidy at the blastocyst stage, aneuploid trophoblast organoids can be readily generated. These models reflect the trophoblast’s unique ability to tolerate aneuploidy and offer a valuable platform to study human placental development and pathophysiology.

## INTRODUCTION

The pre-implantation stage of human embryo development culminates with the formation of a blastocyst, which is ready to implant into the uterus. The blastocyst contains an outer layer of extraembryonic trophoblast cells, a simple epithelial tissue that contributes to the foetal part of the placenta, also known as the chorion, and a small cluster of inner cells known as the inner cell mass (ICM)^1–3^. Upon blastocyst maturation, the ICM segregates into the embryonic epiblast and the extraembryonic hypoblast, which gives rise to the yolk sac. As the mature blastocyst implants into the uterus, the trophoblast initiates the formation of villi, the basic placental unit^4,5^. Primary chorionic villi contain a layer of stem cells, known as villous cytotrophoblasts (CTB) that fuse to form a syncytium, the syncytiotrophoblast (SCT), responsible for gas and nutrient exchange and hormonal regulation^4^. At the tips of the villi, CTB cells proliferate to generate columnar cytotrophoblast (cCTB). These progenitor cells differentiate to form highly migratory cells known as extravillous trophoblast (EVT). EVTs invade deeply within the endometrium and remodel the maternal arteries to increase the blood supply to the placenta^6^. The core of primary chorionic villi is invaded by extraembryonic mesoderm (ExM) cells that play a crucial supportive role, but whose origin remains unknown^7,8^. These ExM cells together with the overlying trophoblast form the secondary chorionic villi.

Defects in early placentation underlie many of the ‘Great Obstetrical Syndromes’, including pre-eclampsia or intra-uterine growth restriction, which have potentially devastating consequences for both the foetus and the mother^9^. However, despite the critical importance of the placenta, our knowledge of the mechanisms that govern its development and function remain very poorly understood. Placentas have experienced constant changes during evolution, with the emergence of different cell types, tissue architectures, and modes of invasion^10,11^. This has limited the use of experimental model systems to understand human placentation. Human trophoblast organoids (TOs) have emerged as a promising tool to study the human trophoblast^12^. These are established from either first trimester placenta^13–15^ or from established cultures of stem cells^16,17^. However, both have limitations. Access to first trimester terminations is becoming increasingly difficult, as these typically happen at home^18^. In contrast, although human embryonic stem cells (ESCs) and trophoblast stem cells (TSCs) are readily available^19,20^, the TOs they form do not contain villous CTB cells but highly proliferative trophoblast stem cells that are prone to undergo EVT differentiation^21,22^. Lastly, current TO models only capture the epithelial trophoblast compartment of the placenta and lack the ExM cells that are an integral part of the chorion. To date, ExM cells have not been derived from human embryos and a model combining both ExM and trophoblast cells to mimic secondary chorionic villi is lacking.

A unique aspect of the human placenta is the relatively high incidence of chromosomal alterations^23^. As embryos develop beyond implantation, aneuploid epiblast cells are selectively eliminated, while aneuploid trophoblast cells persist^24,25^. As a result, in approximately 3% of pregnancies, chorionic villi are trisomic, while the foetus is not affected^26,27^. In many of these cases of confined placental mosaicism (CPM), the placenta is still functional, and healthy babies are born^28,29^. How placental cells cope with the burden of aneuploidy remains unexplored due to the lack of aneuploid *in vitro* models.

Here, we demonstrate that human blastocysts can be used as a starting point to generate TOs with very high efficiency. These TOs recapitulate the major hallmarks of the *in vivo* trophoblast and can be established both from outside and inside cells of the blastocyst. Moreover, given the high incidence of aneuploidy in human blastocysts^24,30^, trisomic TOs can be readily established. Lastly, we have derived the first lines of human ExM cells. We show that these cells originate from inside cells of the blastocyst, and that the hypoblast is needed for their specification. ExM cells and TOs emerge within the same culture conditions and can be combined to model secondary chorionic villi.

## RESULTS

### Derivation of TOs from human blastocysts

To establish TO cultures from human embryos, we initially plated blastocysts on collagen IV / laminin 511-coated plastic plates (2D outgrowth protocol, Figure 1A), similarly to what has been described for TSC derivation^20^. After six days of culture in TSC medium, embryos had attached to the plastic to form an outgrowth reminiscent of TSCs. We passaged the outgrowths into Matrigel domes in TO conditions, where they gave rise to TOs (Figure 1B). We also tried plating the embryos directly in 3D domes of Matrigel in trophoblast organoid medium (3D dome protocol, Figure 1A). After six days in culture, the embryos had collapsed but also grown in size, and upon passaging they generated TO cultures (Figure 1B). Using the 3D dome protocol, the efficiency of TO derivation was approximately 83%, rising to almost 100% when excluding embryos with poor quality at thawing (Table S1).

**Figure 1:**
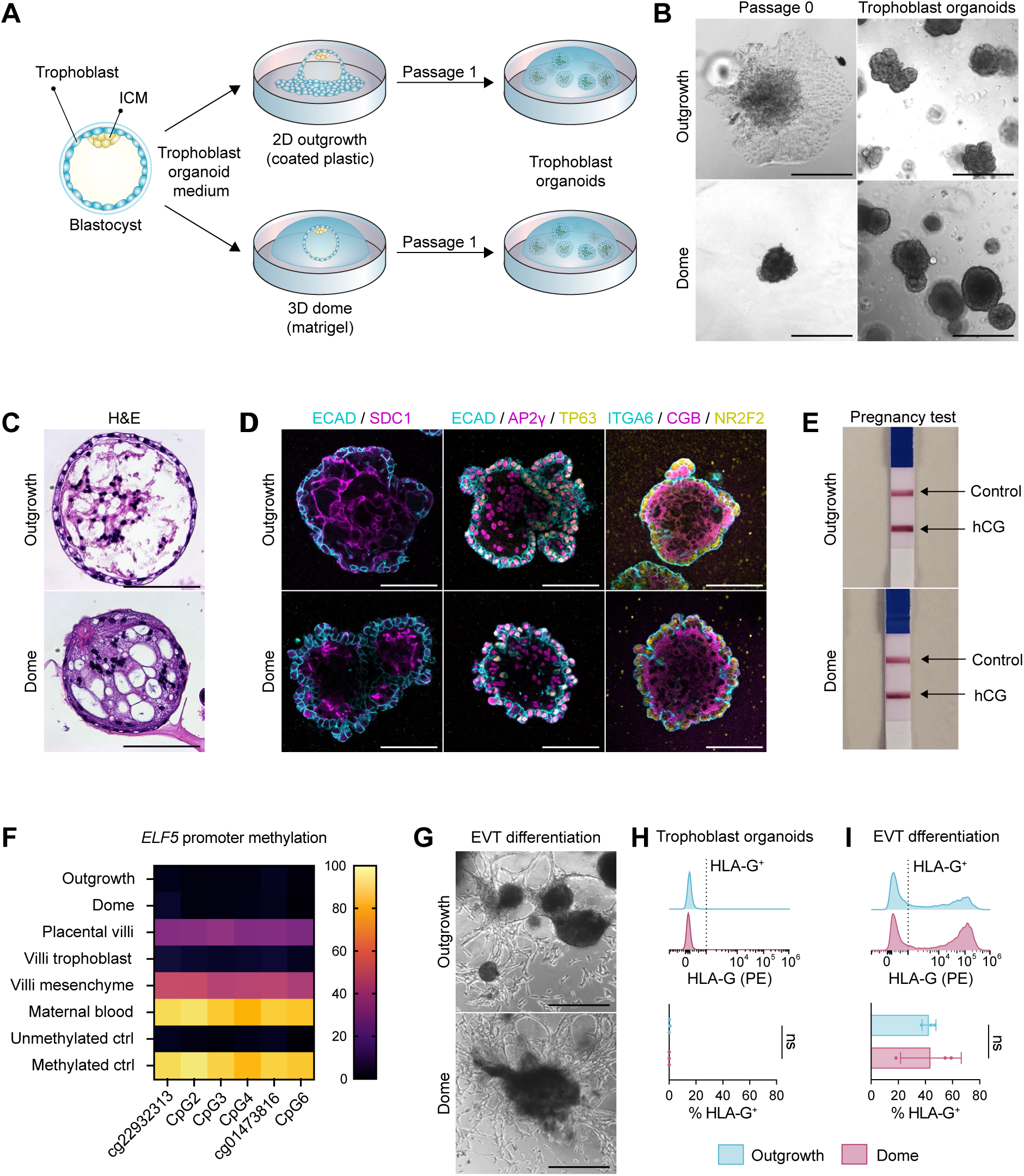
Characterisation of blastocyst TOs. **(A)** Schematic of the derivation protocol. ICM: inner cell mass. **(B)** Phase-contrast images of TO derivations from blastocysts and resulting organoid cultures. Scale bar, 500 µm. **(C)** Haematoxylin and eosin (H&E) staining of TOs derived from blastocysts using two different protocols. Scale bars, 100 µm. **(D)** Immunofluorescence images of TOs. Scale bar, 100 µm. **(E)** Pregnancy test from TO-conditioned media. **(F)** Bisulfite-pyrosequencing of *ELF5* promoter methylation in blastocyst TOs and various tissue samples including relevant controls (ctrl). **(G)** Phase-contrast images of EVT differentiation. Scale bar, 500 µm. **(H, I)** Flow cytometry of Human Leukocyte Antigen (HLA)-G levels of TOs in regenerative conditions and after EVT differentiation. Data is shown as mean ± SEM. n = 3 samples per condition. Two-tailed Student’s t-test, ns: non- significant. 3 independent experiments.

To characterise our blastocyst-derived TOs (herein referred to as ‘blastocyst TOs’) and compare the outgrowth and dome protocols further, we prepared histological sections and carried out wholemount immunofluorescence staining for key trophoblast markers. Both outgrowth and dome TOs exhibited similar gene expression patterns, expressing pan trophoblast markers such as AP2γ, and morphology (Figure 1C). They presented a core of spontaneously differentiated SCT expressing syndecan-1 (SDC1) and human chorionic gonadotropin (CGB) surrounded by an epithelial layer of CTB marked by integrin α6 (ITGA6), E-Cadherin (ECAD), TP63, and NR2F2 (Figure 1D), thus resembling TOs derived from first trimester chorionic villi^13,14^ (from now on referred to as ‘villi TOs’). Blastocyst TOs also secreted CGB into the media, which could be detected with an over-the-counter pregnancy test (Figure 1E) and could be cultured and expanded for more than 6 months. A key feature of trophoblast cells is a distinct methylome, including hypomethylation of the *ELF5* promoter^31,32^. We tested this using bisulphite- pyrosequencing, showing outgrowth and dome TOs exhibit no methylation of *ELF5* promoter CpGs, akin to villous trophoblast from first-trimester pregnancy terminations but distinct from the mesenchymal villous core (Figure 1F).

To determine the capacity of our organoids to make EVT, we differentiated them by withdrawing Wnt agonists from the medium and adding neuregulin-1 (NRG1) as previously described^13^. After two weeks of differentiation, cells with an invasive morphology characteristic of EVT were clearly visible and a human leukocyte antigen (HLA)-G^+^ population was detected by flow cytometry, confirming the capacity of blastocyst TOs to differentiate to an EVT state (Figure 1G-I).

Altogether, our protocol efficiently establishes TOs from human embryos which exhibit hallmarks of *in vivo* placental trophoblast with no observable differences between TOs derived from a 2D outgrowth or directly in a 3D dome.

### Blastocyst TOs are a robust source to establish TSCs

As only a limited number of TSC lines are available^20^, we sought to establish TSC lines from our blastocyst TOs. Following dissociation of TOs, single cells were seeded on collagen IV / laminin-511 coated plates and cultured in TSC medium (Figure S1A). After a few passages, colonies consistently exhibited characteristic TSC morphology, accelerated their growth and could be maintained long term for over 40 passages (Figure S1B). Unlike derivations of TSCs directly from human embryos as described previously^33^, which we found to be inefficient (0/11 embryos,

Table S1), derivation of TSCs from already established TOs was highly efficient. We were able to establish TSC lines from all euploid TOs we tested (n = 9 lines, Table S1).

TO-derived TSCs (from now on referred to as ‘blastocyst TSCs’) showed similar marker expression to Okae TSCs^20^, expressing ECAD, ITGA6 and AP2γ, as well as markers of cCTB such as integrin α2 (ITGA2) and transgelin (TAGLN) (Figure S1C). As with TOs, there was no discernible difference in morphology between TSCs coming from TOs derived using the outgrowth or dome protocol, and both could be differentiated to EVT as previously described^20^ (Figure S1D-S1F). Notably, levels of HLA-G in regenerative conditions were lowest in TSCs originating from dome TOs, intermediate in outgrowth-derived TSCs, and highest in Okae TSCs (Figure S1E). The EVT bias was globally exacerbated in TSCs compared to TOs as previously described^21^, as they have been derived and maintained on 2D plastic dishes.

Overall, our results show that derivation of TSCs via blastocyst TOs is an efficient route for establishing TSCs from human embryos.

### Cell type composition of blastocyst TOs closely resembles *in vivo* placental villi

To compare our blastocyst TOs and TSCs to *in vivo* villous trophoblast and existing TOs, we performed single-cell RNA sequencing (scRNA-seq) of dome and outgrowth blastocyst TOs in regenerative cultures and following EVT differentiation, as well as dome and outgrowth blastocyst TSCs in regenerative cultures. We compared their transcriptomes to *in vivo* first trimester placental villi, villi TOs, and TOs established using Okae TSCs (from now on referred to as ‘Okae TOs’). Following integration of all datasets, we identified 9 cell type clusters and labelled them based on marker gene expression levels (Figure 2A and Table S2). Two CTB clusters (*TEAD4*^+^ / *TP63*^+^ / *EPCAM*^+^) made up the majority of cells *in vivo* and in blastocyst and villi TOs in regenerative medium (Figures 2B). In contrast, a TSC-specific population with similarity to cCTB (*ITGA2*^+^ / *TAGLN*^+^) made up most of the Okae TOs and blastocyst TSCs but was present only at low levels in villi TOs and blastocyst TOs (Figure 2B). Expression patterns of the CTB markers *ELF5*, *BCAM*, *TP63*, *TEAD4* and *EPCAM* in the CTB clusters of blastocyst TOs matched *in vivo* tissue closely, while villi and Okae TOs expressed higher levels of *EPCAM* and reduced levels of *BCAM* (Figure 2C). BCAM has been shown to be characteristic of a primitive CTB progenitor state, while EPCAM is upregulated in maturing CTB^34^. To compare overall similarity between models, we performed Pearson correlation analysis on pseudo-bulk gene expression data per cell-type. Blastocyst TOs closely recapitulated the CTB state found *in vivo,* comparable to villi TOs (Figure 2D). CTB found in Okae TOs and blastocyst TSCs correlated less closely with the *in vivo* tissue.

**Figure 2:**
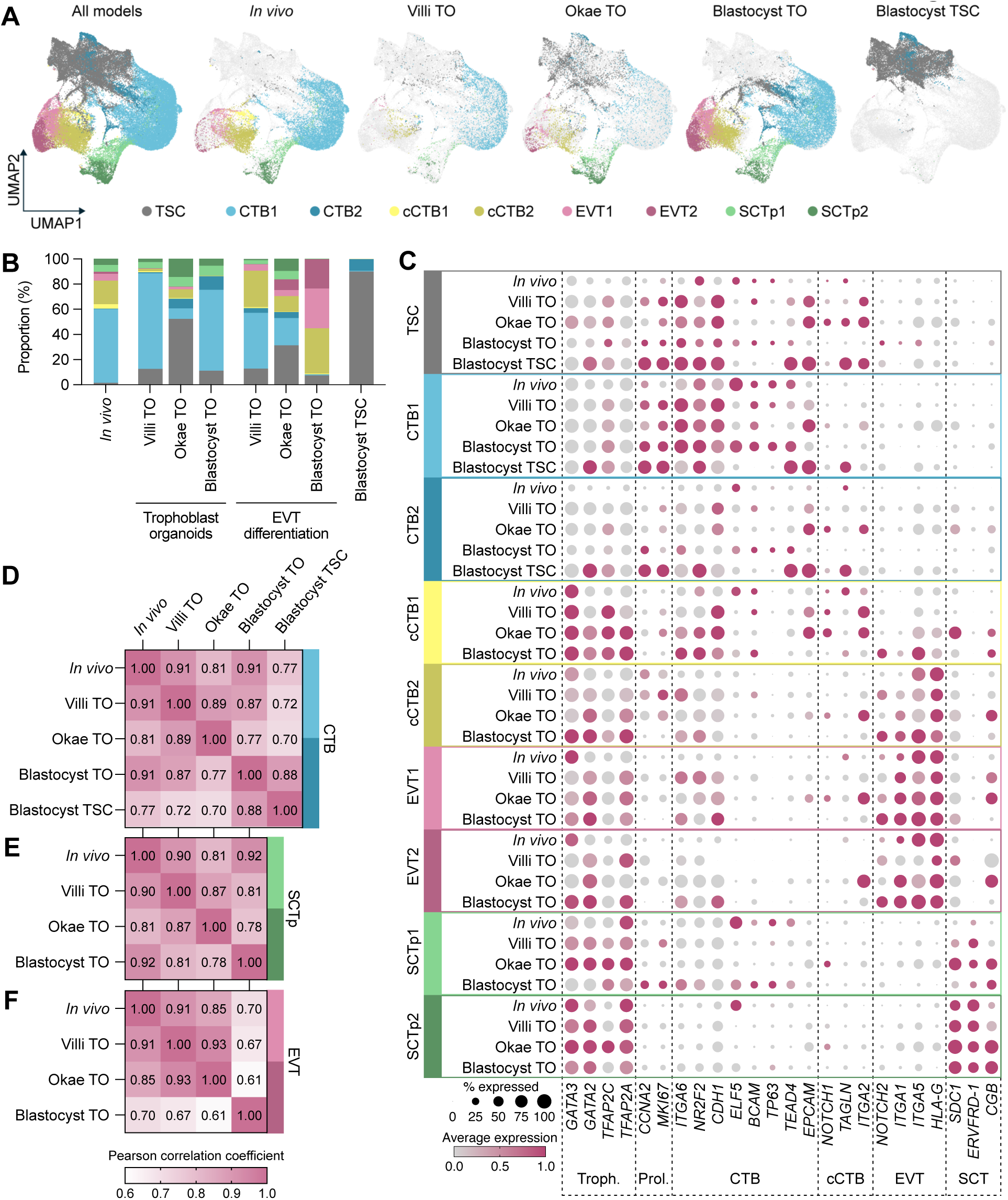
Single cell RNA sequencing analysis of TOs. **(A)** UMAP plot of single- cell transcriptomes of *in vivo* villous trophoblast, villi TOs, Okae TOs, blastocyst TOs, and blastocyst TSCs. TO datasets contain both organoids in regenerative medium as well as after EVT differentiation, while blastocyst TSCs are only from regenerative conditions. **(B)** Proportions of cells in each cluster for each model in A. **(C)** Dot plot of the percentage of cells expressing trophoblast marker genes and the average log expression in each group separated by cluster and trophoblast model. **(D-F)** Heatmap of Pearson correlation analysis comparing the CTB (D), SCTp (E) and EVT (F) clusters between trophoblast models of different origin and the *in vivo* reference.

In terms of the SCT, due to limitations of scRNA-seq, we could only detect SCT precursors (SCTp), while more mature multinucleated SCT was not captured in these datasets. SCTp (*CGB*^+^ / *SDC1*^+^ / *ERVFRD-1*^+^) were identified in all TOs but were near absent from blastocyst TSCs (Figure 2A, 2B) as expected^20^. The expression pattern of marker genes in SCTp from blastocyst TOs matched *in vivo* tissue more closely than villi and Okae TOs, except for increased *CGB* levels (Figure 2C). This was reflected in the Pearson correlation analysis (Figure 2E). In agreement with previous results^22^, we also found the SCT markers CGB and SDC1 to be expressed across different cell types in Okae TOs contrary to their *in vivo* expression pattern (Figure 2C).

EVT differentiation involves the transition of CTB cells into cCTB (*NOTCH1*^+^ / *ITGA2*^+^ / *TAGLN*^+^), which subsequently forms EVT (*ITGA1*^+^ / *HLA-G*^+^)^22^. Upon differentiation of TOs to EVT, the proportion of CTB and SCT decreased while cCTB and EVT cells emerged (Figure 2B). The composition of blastocyst TOs changed most drastically, with very few cCTB and EVT cells in regenerative medium but almost complete differentiation to cCTB and EVT upon EVT differentiation (Figure 2B). However, the EVT state of blastocyst TOs was not as similar as the *in vivo* EVT compared to EVT derived from villi TOs and Okae TOs (Figure 2F). Given that villi TOs and Okae TOs showed more subtle changes in cell composition upon EVT differentiation (Figure 2B), we reasoned the EVT from blastocyst TOs could be at a more mature developmental stage. To explore this further, we clustered the cells from each model separately, retaining the original cell type annotations, and conducted pseudotime analysis with Monocle3 (Figure S2A, S2B). The origin of the trajectory for blastocyst and villi TOs was within the CTB cluster, branching towards either a SCT fate or an EVT fate, reminiscent of the trajectories *in vivo*. However, in blastocyst TOs, the EVT and cCTB clusters were more clearly separated from the remaining cell states, with fewer transitional cell states present and larger pseudotime differences between the CTB origin and the differentiated EVT (Figure S2B). As expected, Okae TOs did not reproduce the *in vivo* differentiation trajectory as differentiated cell types emerged from the TSC cluster (Figure S2B). To corroborate these findings, we looked at additional markers of EVT subtypes and found nearly all of them to be expressed at higher levels in EVT of blastocyst TOs compared to EVT of other TOs or even the *in vivo* reference (Figure S2C).

Lastly, we subset our scRNA-seq data to compare dome and outgrowth TOs, their EVT differentiation, and the TSCs that derive from them. The cell type composition of TOs in regenerative medium and TSCs was highly similar in dome and outgrowth protocols (Figure S2D, S2E). Upon EVT differentiation, the overall composition was comparable as well, although EVT in dome TOs primarily belonged to the EVT2 cluster while outgrowth TOs primarily formed EVT1 (Figure S2E). The expression of EVT genes in blastocyst TOs was similar between the two clusters, besides elevated *CSH1* in EVT2 (Figure S2C).

As we saw differences in basal levels of HLA-G by flow cytometry between dome and outgrowth protocols only in 2D TSCs (Figure S1E), we next compared the expression of trophoblast markers and *HLA* genes between dome and outgrowth TSCs in our scRNA-seq dataset. The pattern of expression between the main TSC cluster and the sparser CTB clusters within each TSC line was highly similar (Figure S2F). Consistent with our flow cytometry data, a small subset of outgrowth TSCs highly expressed *HLA-G*. Outgrowth TSCs also expressed higher levels of cCTB markers (*TAGLN*, *ITGA2),* EVT markers (*ITGA5*), as well as *HLA-A* compared to dome TSCs. These findings support the conclusion that outgrowth TSCs have a more cCTB-like identity than dome TSCs, despite having been cultured under the same conditions for several passages, with the only difference being the protocol for initial TO derivation.

Altogether, single-cell transcriptomics demonstrate that blastocyst TOs closely recapitulate the cell type composition and gene expression patterns of *in vivo* villous placenta, and model some features of trophoblast development more faithfully than TOs from other sources.

### Capturing aneuploidies of human embryos *in vitro* in TOs

Human embryos are frequently aneuploid, with approximately 78% of human blastocysts harbouring aneuploid cells, often being mosaic for different karyotypes^30,35^. While aneuploidy is detrimental in the epiblast and is depleted from this lineage during post-implantation development, the trophoblast tolerates a wide range of trisomies^23,24,36^. To enable modelling of aneuploid trophoblast *in vitro*, we sought to establish aneuploid TOs from human embryos using our protocol. While the majority of the TO lines we derived from intact blastocysts were found to be euploid through karyotyping by shallow whole-genome sequencing (WGS) (10 lines out of 13, Figure S3A and Table S1), 3 lines were found to be aneuploid (Table S1). For example, one line exhibited a trisomy 16 (Figure 3A). While trisomy 16 TOs had similar marker expression to euploid organoids, they formed larger lacunae and took longer to reach confluency than euploid TO lines, consistent with trophoblast hypoproliferation in trisomy 16 embryos^37^ (Figure 3B). Moreover, we could not establish trisomy 16 TSCs from these TOs, unlike any other TO line we tested^37^ (Table S1). As trisomy 16 placentas are associated with adverse pregnancy outcomes^38,39^, these organoids provide a promising platform to study this specific aneuploidy *in vitro*. Next, to assess whether blastocyst TOs acquire *de novo* aneuploidies *in vitro*, we analysed two euploid lines after 8 and 17 passages by shallow WGS, which showed no aneuploidies accumulating at later passages (Figure S3A and S3B). After finding that aneuploid TOs can be established from embryos, we next attempted derivation from embryos that had already been tested as aneuploid before freezing. Pre-implantation genetic testing for aneuploidy (PGT-A) entails the biopsy of 4-5 trophoblast cells of the blastocyst and their analysis by WGS. In six out of seven cases, we found the karyotype of the established TO lines to be discordant with the PGT-A result, such as an embryo diagnosed as monosomy X giving rise to euploid TOs (Figure S3C and Table S1). This highlights the likely presence of euploid cells in the blastocyst and the prevalent mosaicism of human embryos^30^. In one single case, the karyotype of the TOs matched that of the PGT-A biopsy (trisomy 13, Figure S3D). Notably, the efficiency of derivation from aneuploid PGT-A embryos decreased compared to untested embryos (30% for all PGT-A embryos used and 35% when excluding poor-quality embryos) (Table S1). While we were able to establish several TO lines with whole-chromosome trisomies and segmental aneuploidies, derivation was not successful from embryos tested by PGT-A as having autosomal monosomies or more severe complex aneuploidies. The only exception to this was an embryo diagnosed with a monosomy 7 that gave rise to TOs with a partial deletion of chromosome 7 (Table S1). This mirrors the aneuploidies found in cases of CPM^40^, where generally only trisomies are identified, and is consistent with the increased fitness burden of autosomal monosomies and complex aneuploidies^23,41–43^.

**Figure 3:**
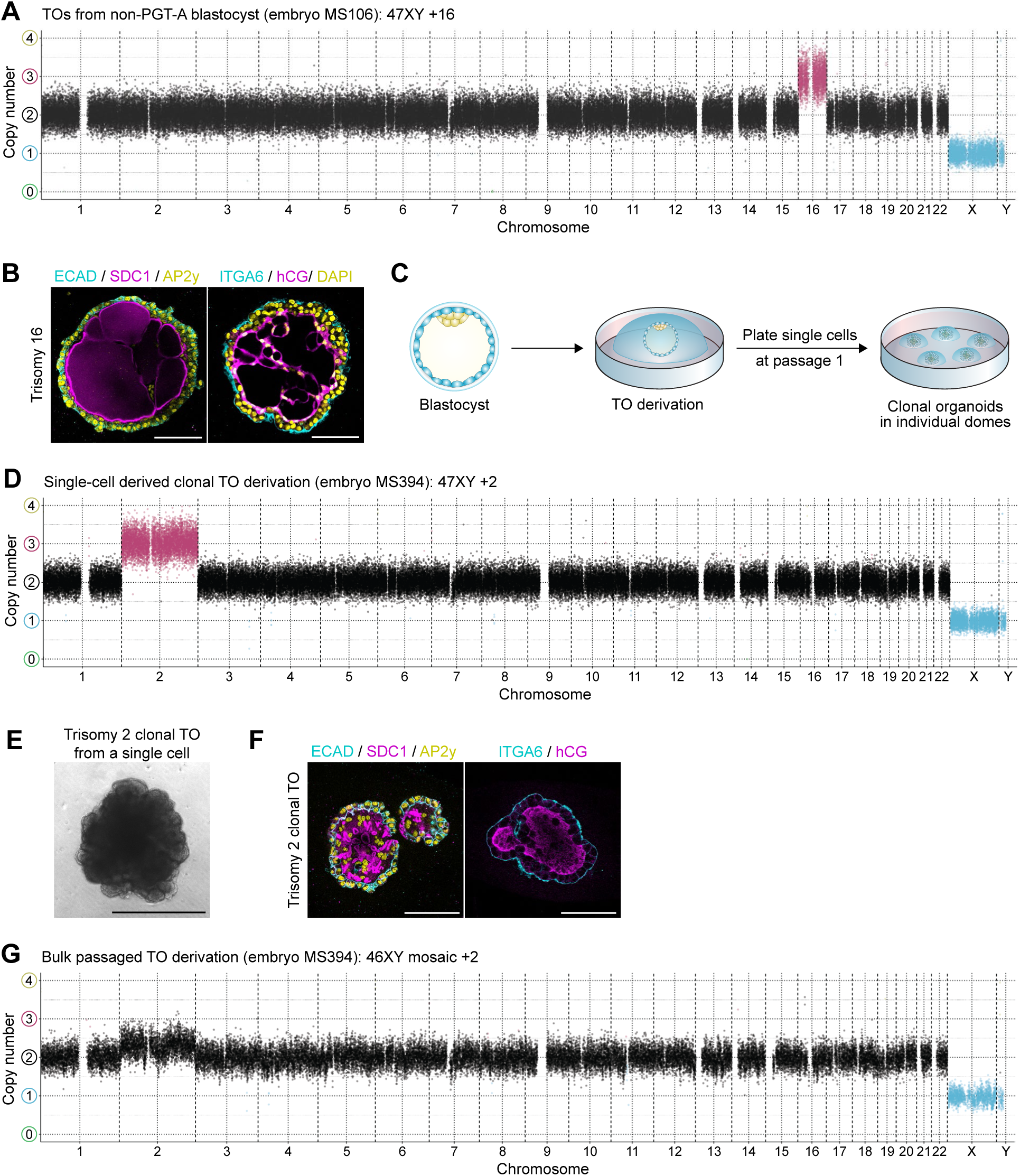
Derivation of aneuploid TOs. **(A)** Copy number analysis of a trisomy 16 TO line. PGT-A: pre-implantation genetic testing for aneuploidy. **(B)** Immunofluorescence images of trisomy 16 TOs. Scale bar, 100 µm. **(C)** Schematic of the clonal derivations. **(D)** Copy number analysis of a trisomy 2 TO line derived from a single cell. **(E)** Phase-contrast image of a trisomy 2 TO grown from a single cell. Scale bar, 500 µm. **(F)** Immunofluorescence images of trisomy 2 TOs. Scale bar, 100 µm. **(G)** Copy number analysis of a mosaic trisomy 2 TO line derived upon passage 1 from the remaining cells of the same embryo as the single cell-derived trisomy 2 TO line.

We speculated that the prevalent mosaicism of human blastocysts may prevent capturing some aneuploidies during derivation. If the embryo contains euploid cells, these could grow faster than the aneuploid cells, leading to dilution and eventual loss of the aneuploid cells from the cultures during passaging. To overcome this limitation, we attempted clonal derivations. Following the initial culture of the embryos using the dome derivation protocol, we manually picked single cells upon passage 1 and plated them in individual domes of Matrigel, while also passaging the remaining bulk (Figure 3C). The efficiency of derivation from single cells was reduced (approximately 20% of single cells giving rise to TOs), but this approach allowed us to obtain clonal lines early during the derivation. Demonstrating the feasibility of this approach to obtain aneuploid TOs from mosaic embryos, we obtained a trisomy 2 TO line from a single cell (Figure 3D-F), while the TOs established from the remaining cells of the embryo were mosaic for trisomy 2 (Figure 3G).

Lastly, we were also able to establish aneuploid TSCs with the karyotype of their parental TOs, which offer a simpler model of aneuploid trophoblast (Figure S3E, S3F and Table S1). The exception to this were trisomy 16 TOs, which we were unable to maintain as a TSC line as previously mentioned. Taken together, our TO derivation protocol can capture trisomies which are naturally occurring in human blastocysts.

### Derivation of TOs from polar and mural halves

Upon blastocyst hatching, trophoblast cells diverge to form two main subtypes, polar trophoblast cells, which are located at the embryonic side, mediate implantation and express markers such as NR2F2, as well as mural trophoblast cells, which are located at the abembryonic side and lack NR2F2 expression^44,45^. To test whether both subtypes have the potential to generate TOs, we dissected blastocysts into mural and polar halves using a laser (Figure 4A-B, Video S1). Both mural and polar pieces could be passaged to establish TO cultures, with the polar half being more efficient at generating TO lines using the dome protocol (8 TO lines from polar halves and 3 TO lines from mural halves; Figure 4C, 4D and Table S3). We observed no apparent differences between mural and polar TOs in marker expression, CGB secretion, and morphology, with mural TOs expressing similar levels of NR2F2 as polar TOs (Figure 4E-H).

**Figure 4:**
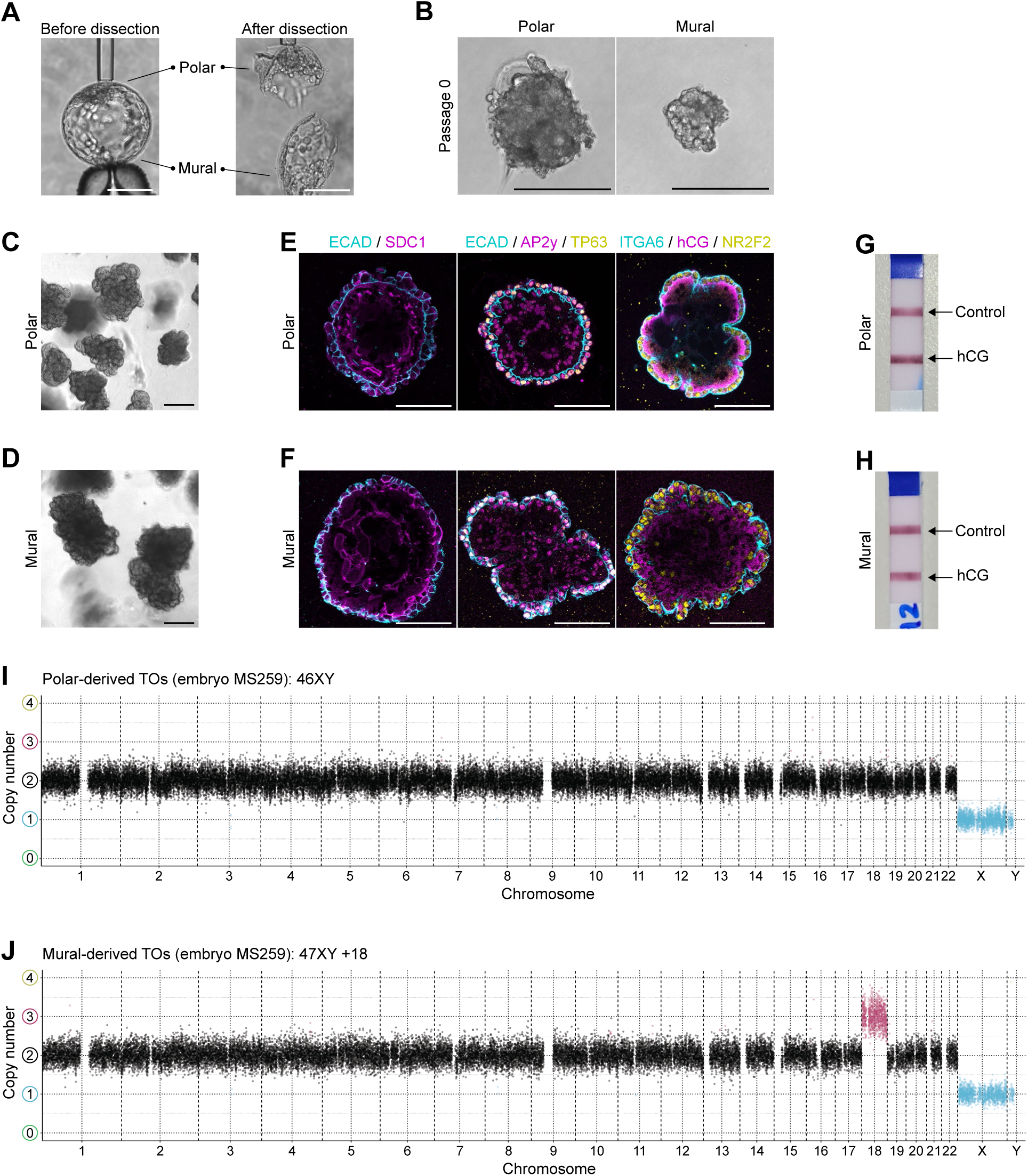
TO derivations from polar and mural halves of the embryo. **(A)** Phase- contrast images showing the laser-assisted dissection of a human embryo into polar and mural halves. Scale bars, 100 µm. **(B)** Phase-contrast images of polar and mural fragments after six days of culture in TO media conditions. Scale bars, 200 µm. **(C, D)** Phase-contrast images of polar and mural TOs. Scale bars, 200 µm. **(E, F)** Immunofluorescence images of polar and mural TOs. Scale bars, 100 µm. **(G, H)** Pregnancy test from conditioned media of polar and mural TOs. **(I, J)** Copy number analysis of TOs derived from the polar and mural halves of a human blastocyst.

The capacity to derive TOs from mural pieces opened the door to deriving isogenic ESCs and TOs from different halves of the same embryo. Euploid ESCs with pluripotency features could be derived from a polar piece following laser dissection (Figure S4A-D), while the remaining mural trophoblast from the same embryo gave rise to TOs (Figure 4D and 4F).

For two embryos, we were able to derive TOs from both their halves (Table S3). Interestingly, in these two cases, the karyotype of the polar and mural TOs was discordant. The polar TOs were euploid in both cases, while the mural TOs exhibited a trisomy 18 as well as a combined trisomy 11 and 13, respectively (Figure 4I-J, S4E-F). This highlights the mosaic karyotypic makeup of human blastocysts. Altogether, our data shows that both mural and polar halves of the embryo can be used to establish TO cultures.

### Derivation of ExM cells from human blastocysts

Having proven that both polar and mural halves can give rise to TOs, we next investigated the fate of ICM cells in our derivations. To do this, we fixed embryos six days after plating in TO conditions, instead of passaging them as usual (Figure 5A). This revealed that while most cells in the embryo expressed the CTB marker ITGA6, a clear cluster of ITGA6^-^ cells was also present (Figure 5B). Only a few ITGA6^-^ cells expressed pluripotency marker genes in approximately 60% of embryos at this stage (Figure 5C). Instead, most of the ITGA6^-^ cells co-expressed the mesenchymal marker vimentin (VIM) and GATA6 (Figure 5D), which is indicative of an ExM identity^7^. We next explored the fate of these cells upon passaging. During the first passages of some TO derivations, we noticed clusters of cells with a distinct mesenchymal morphology alongside TOs (Figure 5E). Immunofluorescence analyses of early passages revealed the presence of cells expressing VIM but lacking expression of trophoblast markers (Figure 5F). We noticed that these putative ExM cells were difficult to passage as they formed a loose pellet that was easily lost. This may explain why ExM cells did not persist beyond the first passages in our previous derivations, as they were aspirated with the supernatant, leaving a stable pellet of mostly trophoblast cells. With an adapted, careful passaging protocol (see Methods), we were able to maintain the ExM cells in culture alongside the TOs.

**Figure 5:**
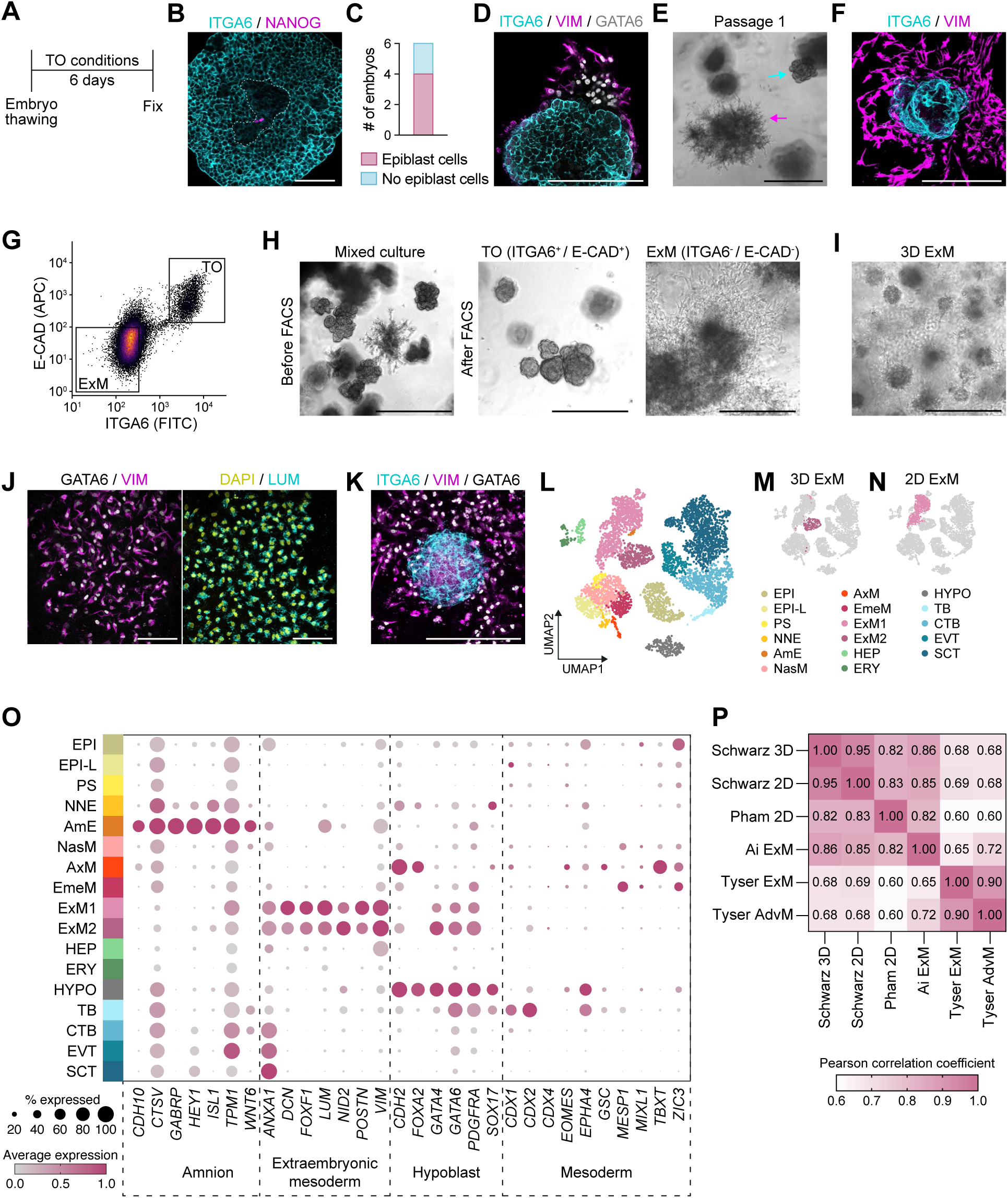
Characterisation of blastocyst ExM cells. **(A)** Schematic of the experiment shown in B-D. **(B)** Immunofluorescence image of a human embryo cultured as indicated in A. Scale bar, 100 µm. **(C)** Contingency bar plot showing the number of embryos from B that have epiblast cells. **(D)** Immunofluorescence image of a human embryo cultured as indicated in A. Scale bar, 100 µm. **(E)** Phase-contrast image of a TO derivation at passage 1. Cells with a mesenchymal morphology (magenta arrow) are present alongside TOs (cyan arrow). Scale bar, 500 µm. **(F) I**mmunofluorescence image (maximum projection) of a TO derivation at early passages. Scale bar, 100 µm. **(G)** FACS result using E-cadherin (E-CAD) and integrin α6 (ITGA6) to sort ExM (E-CAD^-^ / ITGA6^-^) and trophoblast cells (E-CAD^+^ / ITGA6^+^) derived from the same embryo. **(H)** Phase-contrast image of cells before and after separation of ExM and trophoblast cells by FACS. Scale bars, 500 µm. **(I)** Phase-contrast image of blastocyst ExM cultured in 3D. Scale bar, 500 µm. **(J)** Immunofluorescence images of ExM cultured in 3D. Scale bar, 100 µm. **(K) I**mmunofluorescence image (maximum projection) of an ExM-TO co-culture. Scale bar, 200 µm. **(L)** UMAP plot integrating scRNA-seq data from 3D and 2D blastocyst ExM lines with human embryo reference datasets^46–49^. EPI: epiblast; EPI-L: late epiblast; PS: primitive streak; NNE: non-neural ectoderm; AmE: amniotic ectoderm; NasM: nascent mesoderm; AxM: axial mesoderm; EmeM: emergent mesoderm; ExM1: extraembryonic mesoderm 1; ExM2: extraembryonic mesoderm 2; HEP: hemogenic endothelial progenitor; ERY: erythrocyte; HYPO: hypoblast; TB: pre- implantation trophoblast; CTB: cytotrophoblast; EVT: extravillous trophoblast; SCT: syncytiotrophoblast;. **(M)** 3D ExM highlighted on the UMAP plot from L. **(N)** 2D ExM highlighted on the UMAP plot from L. **(O)** Dot plot of the percentage of cells expressing amnion, ExM, hypoblast and mesoderm marker genes and the average log expression in each cluster of the UMAP in L. **(P)** Heatmap of Pearson correlation analysis comparing 2D and 3D blastocyst ExM (‘Schwarz’), ESC-derived 2D ExM (‘Pham’)^7^, ExM from *in vitro* cultured human embryos (‘Ai’)^47^ and advanced mesoderm (AdvM) and ExM from an *in vivo* gastrulating embryo (‘Tyser’)^49^.

To resolve the heterogeneity of the ExM-TO cultures, we sorted cells using the trophoblast surface markers ITGA6 and E-CAD (Figure 5G). This allowed us to obtain homogeneous 3D cultures of double-positive TOs and double-negative ExM cells (from now on referred to as ‘blastocyst ExM’) (Figure 5H). Under these conditions, blastocyst ExM cells formed 3D aggregates, with some mesenchymal cells invading the Matrigel as either single cells or groups of cells (Figure 5I). Unlike TOs, 3D ExM cells did not secrete CGB (Figure S5A) and expressed the ExM markers GATA6, VIM, and lumican (LUM) (Figure 5J). Moreover, ExM cells lacked E-CAD expression and instead expressed N-cadherin (N-CAD) (Figure S5B).

ExM cells emerge when pre-implantation-like human ESCs are exposed to TSC medium^7^. Since these ESC-derived ExM cells are expanded in 2D^7^, we tested whether blastocyst ExM cells could be expanded in 2D as well. We seeded the cells on collagen IV-coated plates in 2D ExM medium^7^ and obtained ExM cells with a morphology and a marker expression similar to ExM cells differentiated from ESCs (Figure S5C, S5D). Karyotyping by shallow WGS revealed that 5 out of 7 blastocyst ExM lines were euploid (Figure S5E and Table S3), but we also identified a trisomy 22 line and a trisomy 8 with an additional segmental duplication of chromosome 8 and a small deletion on chromosome 12 (Figure S5F, S5G). In two cases where we were able to derive both TOs and ExM from the same embryo, the two cell types had a different karyotype (Figure S5G, S5H, Table S3), once more highlighting the mosaic karyotypic makeup of human blastocysts.

Current TO models do not incorporate the mesenchymal component that is characteristic of secondary chorionic villi. To explore if ExM lines could be combined with TOs from other embryos, we mixed both cell types together within a 3D Matrigel dome. The two cell types could be co-cultured, and ExM cells were in contact with CTB cells, mimicking the spatial arrangement of cell types in chorionic villi as they develop from primary villi to secondary villi^46^ (Figure 5K). Therefore, ExM and TOs can be combined to study their interactions and maturation of chorionic villi.

### Transcriptional characterisation of blastocyst ExM cells

For further characterisation of our blastocyst ExM lines, we carried out scRNA-seq of ExM cultures in 3D as well as in 2D and integrated these data with scRNA-seq datasets from human pre- and post-implantation embryos^47–50^ (Figure S6A). After clustering, we annotated embryonic cell types based on marker expression resulting in 17 clusters (Figure 5L and Table S4). Comparison to the original annotations of the individual embryo datasets showed overall agreement with our cell type annotations (Figure S6A). Blastocyst ExM cultured in 3D and 2D belonged mainly to the ExM2 and ExM1 clusters respectively (Figure 5M, 5N). Both ExM clusters highly expressed a characteristic panel of markers including *VIM*, *LUM*, *POSTN, FOXF1*, as well as *GATA4*, *GATA6,* and *PDGFRA* (Figure 5O and Table S5), consistent with immunofluorescence images of blastocyst ExM (Figure 5J, S5D). They did not express markers specific to embryonic mesoderm such as *TBXT*, *ZIC3,* and *MESP1* (Figure 5O). Of the embryo reference cells, cells annotated as ‘advanced mesoderm’ and ‘extraembryonic mesoderm’ by Tyser and colleagues^50^ were located within the ExM1 cluster correlating to our 2D ExM, while ExM cells from the Ai dataset were found mainly in the ExM2 cluster of the integrated UMAP together with our 3D ExM (Figures S6A, S6B).

Pseudobulk comparison of blastocyst ExM with 2D ExM cells differentiated from human ESCs^7^ (‘Pham 2D’) showed high similarity, especially between cells cultured in 2D in the same conditions (Figure 5P). When compared to the original clusters from the Tyser^50^ and Ai^48^ datasets, blastocyst ExM correlated better with the embryo references than ExM derived from ESCs (Figure 5P).

As another way to compare blastocyst ExM to available human embryo scRNA-seq datasets, we used a published early embryogenesis prediction tool^51^. Consistent with our own integrated dataset, 2D and 3D blastocyst ExM mapped to ExM from available embryo reference datasets (Figure S6D-F). Overall, our analysis demonstrates that the transcriptome of blastocyst ExM closely resembles ExM from available human embryo reference datasets, enabling the *in vitro* study of this mesenchymal compartment.

### Tissue requirements for ExM cell specification

We next sought to investigate the tissue requirements for ExM specification. First, we dissected embryos into polar and mural halves and fixed them after six days of culture in TO conditions (Figure 6A). This revealed that ExM cells emerged exclusively from the polar half (Figure 6B, 6C). Since the polar region of the embryo contains both trophoblast and the ICM, we next performed immunosurgery on intact blastocysts to kill outside trophoblast cells and test the capacity of inner cells to make ExM (Figure 6D). After six days of culture in TO conditions, we noticed some ICMs resembled TOs in morphology, while others had cells with a clear mesenchymal morphology (Figure 6E). By fixing the ICMs at this point in the protocol, we also observed VIM^+^/GATA6^+^ ExM cells alongside ITGA6^+^/NR2F2^+^ trophoblast cells coming from the same ICM (Figure 6F). Depending on the ICM, passaging gave rise to ExM, TOs or both (Table S3). These results indicate that the ICM retains a high level of plasticity, and under the right culture conditions it can give rise to both trophoblast and ExM cells.

**Figure 6:**
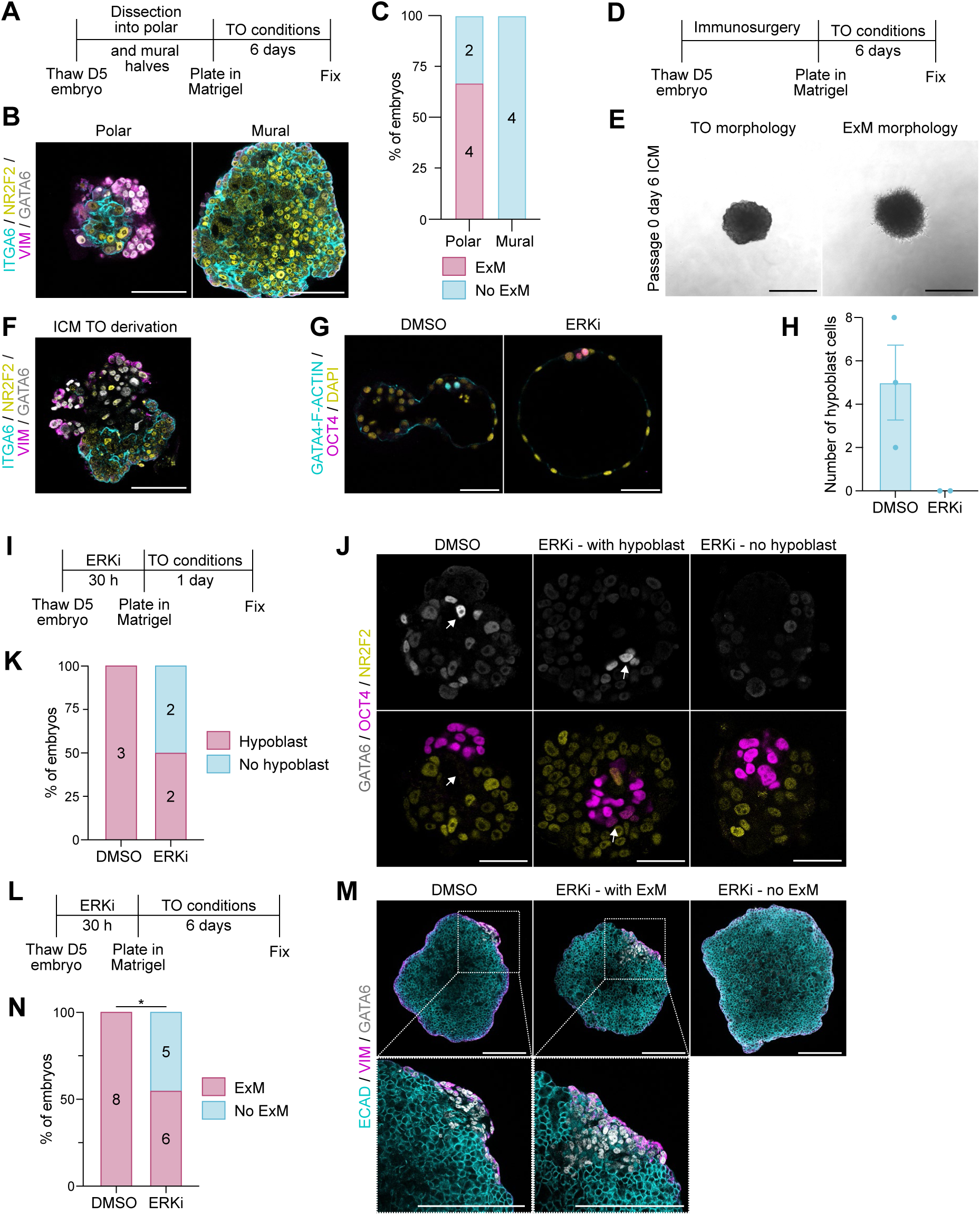
The hypoblast is needed for ExM cell specification. **(A)** Schematic of the experiment shown in B-C. **(B)** Immunofluorescence image of polar and mural halves fixed after six days of culture in TO conditions. Scale bar, 100 µm. **(C)** Contingency bar plot showing the number of embryos from B that have ExM cells. **(D)** Schematic of the experiment shown in E-F. **(E)** Phase-contrast images of ICMs after six days of culture in TO conditions. Scale bars, 500 µm. **(F)** Immunofluorescence image of an isolated ICM fixed after six days of culture in TO conditions. Scale bar, 100 µm. **(G)** Immunofluorescence images of embryos after 30 hours of DMSO or ERK inhibitor treatment. Scale bars, 50 µm. **(H)** Number of hypoblast cells in embryos from G. Data is shown as mean ± SEM. Each dot represents an individual embryo. 3 independent experiments. **(I)** Schematic of the experiment shown in J-K. **(J)** Immunofluorescence images of embryos cultured as specified in I. Scale bars, 50 µm. **(K)** Contingency bar plot showing the number of embryos from J that have hypoblast cells. 2 independent experiments. **(L)** Schematic of the experiment shown in M-N. **(M)** Immunofluorescence images of embryos cultured as specified in L. Scale bars, 200 µm. **(N)** Contingency bar plot showing the number of embryos from M that have ExM cells. Two-sided Fisher’s exact test, *p ≤ 0.05. 5 independent experiments. ERKi: ERK inhibitor.

Since the ICM gives rise to both epiblast and hypoblast cells, both of which have been proposed as the cell type of origin for the ExM^8^, we next explored the role of the hypoblast during ExM specification. To understand whether the hypoblast is needed for ExM formation, we inhibited hypoblast formation in day 5 blastocysts by subjecting them to ERK inhibition for 30 hours as previously described^52^. We could confirm that ERK inhibition completely blocked hypoblast specification (Figure 6G, 6H). Following this pre-treatment, we plated the embryos in TO conditions. We first fixed the embryos after one day (Figure 6I) to analyse whether removal of the ERK inhibitor and culture in TO medium would allow the re-specification of hypoblast cells. We observed that while all DMSO control embryos had GATA6^+^/OCT4^-^ cells, 50% of embryos that had been pre-treated with the ERK inhibitor lacked GATA6^+^ cells (Figure 6J, 6K). Interestingly, the remaining 50% had GATA6^+^/OCT4^+^ cells (Figure 6J, 6K). As OCT4 is expressed in the early hypoblast but becomes downregulated as hypoblast cells mature^53^, this result suggests that the hypoblast cells present in 50% of the embryos pre-treated with the ERK inhibitor emerge during the derivation protocol, once ERK inhibition is relieved. Next, we pre-treated the embryos with the ERK inhibitor for 30 hours and cultured them in TO conditions for six days to assess ExM specification (Figure 6L). ERK inhibition at the blastocyst stage significantly impacted the formation of ExM cells compared to the DMSO group, with around half of the embryos unable to form ExM (Figure 6M, 6N). Importantly, the ratio of embryos pre-treated with the ERK inhibitor and still able to form ExM was consistent with the ratio of embryos re-forming hypoblast during the first day of culture under the same conditions. Overall, these data suggest that the hypoblast plays an important role in ExM formation during derivation from human embryos.

## DISCUSSION

A healthy placenta is absolutely required for a healthy baby to be born. And yet, our current knowledge of human placental biology is very limited due to technical challenges when studying this organ *in vitro*. We have developed a robust and reproducible protocol to derive human trophoblast organoids from early human embryos. Despite their early embryonic origin, the resulting organoids are transcriptionally and morphologically similar to organoids derived from first trimester placentas of pregnancy terminations. This indicates the culture conditions used promote the maturation of the early trophoblast lineage of the blastocyst.

By using blastocysts as a starting point of the derivations, we have been able to investigate key aspects of early human embryo development. We have explored the plasticity of the cells in the human blastocyst. Our experiments revealed that inner cells of the blastocyst retain the ability to form trophoblast, in agreement with previous findings^54^. Which type of inside cells give rise to trophoblast remains uncertain, and several possibilities can be hypothesised. A recent report has demonstrated the existence of inside GATA3^+^ cells in human blastocysts, which increase in number as blastocysts are cultured *in vitro*^55^. The fate of these cells is currently unknown, but since they express trophoblast markers it is tempting to speculate that they are the cell type of origin of the TOs. In mouse embryos, the hypoblast has been found to retain multi-lineage plasticity^56^. Whether this is also the case in human embryos has not been explored, and therefore we cannot rule out the possibility that hypoblast cells contribute to TO development. Lastly, cells that express the naive pluripotency marker SUSD2 have been shown to form GATA3^+^ cells *in vitro*^54^, and therefore the epiblast could potentially also give rise to TOs.

Our experiments have shown that when blastocyst or specifically inside cells are cultured in TO conditions, not only trophoblast cells develop but ExM cells grow alongside. This is reminiscent of the *in vitro* specification of ExM from human ESCs under TSC conditions^7^. Importantly, we have separated ExM cells from trophoblast cells and generated the first lines of ExM cells derived from human embryos. These cells can readily grow both in 2D and 3D conditions, and under these two conditions they capture different ExM sub-populations of human embryos based on their transcriptional profile. The blastocyst ExM cells can also be combined with TOs to form a model of secondary chorionic villi that includes both the epithelial and mesenchymal compartments of the foetal placenta. This model represents a powerful *in vitro* tool to dissect the understudied crosstalk between ExM and trophoblast cells.

The cell type of origin and tissue requirements for ExM specification in human embryos remains unknown. Most of our current knowledge is derived from histological examinations of *in vivo* developing human and non-human primate embryos^57,58^ or inference from scRNA-seq data of embryos and stem cell-based embryo models^7,48^. Critically, different stem cell-based embryo models disagree on the origin of human ExM^3^, highlighting the importance of addressing this question directly in embryos. Our experiments have shown that mural trophoblast cells do not give rise to ExM under our experimental conditions, indicating a restriction in their developmental potential compared to inner cells of the blastocyst. To dissect the contribution of epiblast and hypoblast cells to ExM specification, we investigated the appearance of ExM cells in embryos in which hypoblast specification was inhibited. This revealed a clear decrease in the efficiency of ExM specification, indicating the hypoblast contributes to the formation of ExM in human embryos. Whether this reflects a direct or an indirect role remains unknown. Future lineage tracing experiments will be needed to shed light on this question.

A unique aspect of the trophoblast is the capability to tolerate aneuploidy^25,26^. Aneuploidy is universally detrimental for cellular physiology^59^ and yet, healthy placentas frequently display chromosomal aberrations, specifically trisomies^23,27^. How placental cells cope with the burden of aneuploidy remains unknown due to the lack of *in vitro* models of the aneuploid placenta. Here, we have leveraged the high incidence of aneuploidy in human blastocysts^30,60^ to derive aneuploid TOs. While we successfully derived a battery of trisomic TOs, we failed to establish whole chromosome monosomic and complex abnormal TOs. This mimics observations from chorion villous sampling, in which trisomies spanning all autosomes but chromosome 1 can be detected, but monosomies or more complex karyotypes are typically not found^40^. Our collection of trisomic TOs provides a unique opportunity to dissect chromosome-specific effects and general responses of the trophoblast to aneuploidy. For example, trisomy 16 CPM is of clinical significance, as it is associated with placental defects^61^, which manifest shortly after implantation^37^. However, the lack of *in vitro* models has hampered the identification of underlying mechanisms. We have established trisomy 16 TOs which recapitulate the trophoblast phenotype of trisomy 16 embryos and will allow future studies into the molecular mechanisms behind placental dysfunction.

Our derivations have also highlighted the high degree of mosaicism of human blastocysts and the lack of accuracy of PGT-A as a diagnostic tool. Out of seven embryos diagnosed as aneuploid by PGT-A, TOs with a matching karyotype were obtained only in one case. Moreover, the dissections of human embryos into polar and mural halves consistently gave rise to TOs with different karyotypes. We also obtained TOs and ExM with different karyotypes from the same embryo. This result reflects the different types of CPM, namely CPM type I, in which aneuploid cells are present in the trophoblast but not mesenchymal core, CPM type II, in which aneuploid cells are restricted to the mesenchymal core, and CPM type III, which affects both tissues^40^. Future studies using our model of secondary chorionic villi and

TOs and ExM of different karyotypes will allow us to investigate these different modalities of CPM. Finally, given the increasing numbers of supernumerary human embryos being generated through medically assisted reproduction, we anticipate this methodology could be used to study placental function in a personalised manner in the future.

## Limitations of the Study

Although we have demonstrated that euploid TOs are genetically stable, we do not know whether trisomic TOs display genomic instability and accumulate additional chromosomal alterations upon prolonged *in vitro* culture. Along similar lines, the lack of concordance between the PGT-A result and the karyotype of the TOs, could be due to the accumulation of additional chromosomal alterations upon *in vitro* culture and/or the activation of correction mechanisms.

Regarding the ExM, the current embryo references to date do not capture the maturation of ExM into more mature cell types of the tertiary chorionic villi, limiting a more granular analysis of ExM subtypes and their differentiation trajectories. Additionally, they lack a spatial component. Therefore, we do not know whether our ExM cells match more closely the chorionic ExM or ExM from other extraembryonic organs, such as the amnion or the yolk sac, and indeed if and how these spatially separate populations of ExM differ *in vivo*. trajThe co-culture of ExM and TOs recapitulates the spatial arrangement of cell types in the secondary chorionic villi, but the SCT is inside the organoid, rather than outside, as in canonical TO models^13,14^. Future experiments combining inversion protocols for TOs^62–64^ together with ExM cells will be needed to improve the model.

Lastly, although our experiments suggest the hypoblast plays a role during ExM specification, we cannot determine the cell type of origin of the ExM due to the lack of lineage tracing tools in human embryos.

## ACKNOWLEDGMENTS

We thank Magdalena Zernicka-Goetz, Kathy Niakan, Thorsten Boroviak, Nanami Satoh, and Miguel Angel Ortiz Salazar for their valuable feedback. We are grateful to Zoe Barnikel, Catherine Pretty, Alex Page, Caroline Drew, Sammantha Thodes, Hannah Newby, Yvonne Lodge, Kathryn Berrisford, Lynne Nice, Katie Davies, Bethany Marshall, Ebele Iloabachie, Laura Rhodes, Megan Lockwood, Sandra Gutierrez, and Su Barlow from CARE Fertility clinics for their help organising human embryo donations for research. We thank Angel Martin Bastida for his help with embryo thawing. L.C.S. is funded by the Gates Cambridge Trust. R.C.S. is funded by the Funai Foundation for Information Technology. R.V-T. lab’s is funded by UK Research and Innovation (UKRI) under the UK government’s Horizon Europe funding Guarantee (grant number EP/Y009924/1) and the Wellcome Trust (grant 220540/Z/20/A). Work done in A.B.’s lab is funded by the Natural Sciences and Engineering Research Council of Canada Discovery (RGPIN-2020-05378) and the Canadian Institutes of Health Research Project Grant (202109PAV-468535-CA2). The laboratory of M.N.S. is funded by the Medical Research Council as part of UK Research and Innovation (grant reference MRC, MC_UP_1201/24), the Engineering and Physical Sciences Research Council (Horizon Europe guarantee funding EP/X023044/1) and the EMBO young investigator programme.

## AUTHOR CONTRIBUTIONS

L.C.S. designed, performed, and analysed most of the experiments. M.J.S. analysed the scRNA-seq data from TOs and TSCs. G.M. performed basic characterisations of the TOs. R.C.S. analysed the scRNA-seq data from ExM cells. V.S.R. helped with human embryo thawing and manipulation. K.C. performed the ELF5 promoter methylation experiment. L.K. performed human ESC derivations and characterised the resulting human ESCs. P.S., L.C., K.E., A.M., L.W., R.G., T.B., I.S., A.B., and A.C. provided human embryos for this study. R. V-T. secured funding for the scRNA-seq of TOs and TSCs. A.G.B. oversaw the characterisation of TOs and the scRNA-seq analyses of TOs and TSCs. M.N.S. designed, performed, and analysed embryo experiments, supervised and conceived the project. The manuscript was written by L.C.S. and M.N.S. with help from co-authors.

## DECLARATION OF INTERESTS

The authors declare no competing interests.

## METHODS

### Ethics statement

Most of the human embryo experiments were performed at the MRC Laboratory of Molecular Biology. This experimental work was done in accordance with Human Fertility and Embryology Authority (HFEA) regulations (license reference R0207). Ethical approval was obtained from the National Health Service (NHS) Research Ethics Committee (REC) Cambridgeshire and Hertfordshire (REC reference 19/EE/0381). Informed consent was obtained from all the patients that donated their supernumerary and cryopreserved human embryos from Bourn Hall clinic, CARE Fertility clinics, and King’s Fertility clinic. Prior to giving consent, patients were informed about the specific objectives of the project and the conditions of the license. Patients did not receive any financial inducements, and they were offered counselling.

Some of the dissections of human embryos into polar and mural sides were performed at the University of Michigan. These experiments were approved by an Institutional Review Board (study HUM00028742). Prior to giving consent, patients were informed about the specific objective of the project and the conditions of the donation. Patients did not receive any financial inducements, and they were offered counselling. Following human ESC production and characterization, documents demonstrating adherence to NIH-established guidelines for embryo donation and human ESC production were submitted to NIH for placement on the NIH human ESC Registry and approval was granted on 02/02/2012 (NIHhESC-12-0147). Derivation of human ESCs and their derivatives were performed with non-federal funds.

### Human embryo thawing

Cryopreserved day 5 or day 6 human embryos were thawed using Kitazato thaw kit (VT802- 2, Hunter Scientific) following the manufacturer’s instructions. Briefly, an individual straw containing a donated human embryo was taken from liquid nitrogen and immediately placed in TS solution for 1 minute. This was followed by a 3-minute incubation in DS, 5-minute incubation in WS1, and 1 minute incubation in WS2 in reproplates (REPROPLATE, Hunter Scientific). Embryos were then washed through 3 drops of Global Total LP human embryo culture medium (H5GT-010, LifeGlobal group) and cultured for 2-4 hours in drops of Global Total LP under mineral oil (9305, Fujifilm Irvine Scientific) to allow recovery from thawing and expansion. Global Total LP medium was equilibrated overnight at 37 °C in 21% O_2_ and 5% CO_2_ prior to embryo thawing. Human embryos were cultured at 37 °C in 21% O_2_ and 5% CO_2_. After recovery, embryo quality was assessed, and embryos were assigned a specific grade as follows: 1. very poor quality, collapsed embryo with several dead cells; 2. poor quality, collapsed embryo with a few dead cells; 3. intermediate quality, small cavity present with a few dead cells; 4. good quality, expanded blastocyst with TE and ICM present; 5. very good quality, expanded blastocyst with a well-developed TE and ICM.

### Human embryo manipulations and treatments

The following manipulations to the human embryos were performed:

- Polar-mural dissections: fully expanded day 6 human embryos were placed in a drop of pre-warmed G-PGD (10074, Vitrolife) supplemented with 5 mg/mL HSA-solution (10064, Vitrolife). The dissection was done using a Zeiss Axio Observer D1 microscope coupled with Eppendorf injectMan 4 micromanipulators. The embryo was hold in place through the polar TE using a VacuTip II holding capillary (5195000.044, Eppendorf). A biopsy micropipette (MBB-FP-SM-30, Cooper Surgical) was used to pull on the mural TE. 4.5 millisecond laser shots (Octax, Vitrolife) were used to cut the blastocyst perpendicular to the polar-mural axis (Video S1). In a few cases, the laser was not sufficient to fully separate the two sides, and a finely pulled glass capillary was used to separate the polar and mural halves. These were plated in growth factor-reduced Matrigel (356231, Corning) and cultured as described for “dome” derivations below. Mural TE pieces generated in the University of Michigan were cryopreserved by vitrification. Briefly, the TE biopsies were incubated for 1 minute in pre-warmed solution A at 37 °C. This was followed by a 25 second incubation in a drop of pre-warmed solution B at 37 °C, and an additional wash with a fresh drop of pre-warmed solution B. The TE biopsies were then loaded onto the cryolock with minimal solution and immediately plunged into a liquid nitrogen cooler. Solution A contains 10% DMSO and 10% ethylene glycol in G-MOPS Plus (10130, Vitrolife) and solution B contains 205 DMSO, 20% ethylene glycol, and 0.3 M trehalose in G-MOPS Plus.
- Immunosurgery: the trophoblast of day 6 human embryos was removed by immunosurgery as previously described^65^ with small modifications. Briefly, the zona pellucida of fully expanded day 6 human embryos was first removed by a short exposure to Acidic Tyrode’s solution (T1788, Sigma), and then the embryos were cultured in a 1/5 dilution of anti-human antiserum (H3383, Sigma) diluted in N2B27 for 45 minutes at 37 °C. The embryos were subsequently washed three times with N2B27 and incubated for 45 minutes in a 1/5 dilution of rat serum (provided by Charles River laboratories). At this point, the swelling of the TE became evident, and the embryos were incubated for an additional 45 minutes in N2B27. The lysed TE cells were removed using a finely pulled glass capillary, and the isolated ICMs were either fixed immediately or plated in TO conditions as described below.
- ERK inhibition: day 5 human embryos were thawed and immediately cultured in Global Total LP with either 5 µM Ulixertinib or DMSO (control) for approximately 30 hours in ultra-low attachment 96-well U-bottom plates (7007, Corning) without mineral oil. At that point, the embryos were plated in TO derivation conditions as described below and fixed six days after plating.

### TO derivations

Following thawing and recovery, human blastocysts were treated with Acidic Tyrode’s solution (T1788, Sigma) to remove the zona pellucida. For “outgrowth” derivations, 4-well plates were coated with 5 µg/mL collagen IV (354233, Corning) and 0.5 µg/mL iMatrix-511 (T303, Takara Bio) in PBS for 3 hours at 37 °C. Blastocysts were plated and cultured in TSC medium, with a media change after three days. After six days, the blastocyst outgrowth was dissociated with TrypLE Express (12604021, Gibco) for 12 min at 37 °C. Single cells and small aggregates of cells were obtained by manual pipetting with a 175 µm stripper trip (MXL3-175, CooperSurgical) followed by a 75 µm stripper tip (MXL3-75, CooperSurgical) in wash buffer. The cell suspension was then added to a 40 µL drop of growth factor-reduced Matrigel (356231, Corning) in a 24-well plate. Matrigel was allowed to solidify for 2 min at 37 °C before turning the plate upside down to distribute cells within the drop for 30 min at 37 °C. For “dome” derivations, blastocysts were plated in individual 40 µL dome of growth factor-reduced Matrigel per well in a 4-well dish and the plate was turned over for 30 min at 37 °C as described above to prevent blastocysts attaching to the bottom of the dish. 650 µL TOM was added to the wells and replaced after three days. On day six of the derivation, TOM was removed and 1 mL of Dispase (07923, STEMCELL Technologies) was added to the well. The Matrigel dome was then broken by manual pipetting and the Matrigel-Dispase mix was incubated at 37 °C for 20 minutes. The embryo was then placed on a drop of TrypLE Express for 12 min at 37 °C and a cell suspension was obtained and plated in a drop of growth factor-reduced Matrigel as described for the “outgrowth” protocol.

TOM contains Advanced DMEM/F12 (12634010, Thermo Fisher Scientific), 1X B27 (12587010, Gibco), 1X N2 (homemade), 1X GlutaMAX (35050061, Thermo Fisher Scientific), 100 µg/mL Primocin (ant-pm-05, Invivogen), 1.25 mM N-acetyl-L-cysteine (A9165, Sigma-Aldrich), 100 ng/mL zebrafish bFGF (Marko Hyvönen lab, University of Cambridge), 80 ng/mL human R-spondin 1 (120-38-100ug, PeproTech), 50 ng/mL human EGF (Qk011, Qkine), 50 ng/mL human HGF (100-39H-100ug, PeproTech), 2.5 µM Prostaglandin E2 (P0409, Sigma-Aldrich), 2 µM Y27632 (Y-27632, STEMCELL Technologies), 1.5 µM CHIR99021 (72054, STEMCELL Technologies) and 500 nM A83-01 (72022, STEMCELL Technologies). N2 supplement contains DMEM/F12 (21331-020, Thermo Fisher Scientific), 0.75% bovine albumin fraction V (15260037, Thermo Fisher Scientific), 2.5 mg/ml insulin (I9287, Sigma-Aldrich), 10 mg/ml Apotransferrin (T1147, Sigma-Aldrich), 2 mg/ml progesterone (p8783, Sigma-Aldrich), 0.6 mg/ml sodium selenite (S5261, Sigma-Aldrich) and 1.6 mg/ml putrescine dihydrochloride (P5780, Sigma-Aldrich). Wash buffer contains DMEM/F12 and 0.1% BSA (A3311, Sigma-Aldrich).

### TO culture, maintenance, and passaging

TOs were cultured at 37 °C in 21% O_2_ and 5% CO_2_ in TOM. Fresh medium was added every two to three days. When organoids reached a size of ∼500 µm they were released from Matrigel domes with Dispase for 20 min at 37 °C and mechanical disruption by pipetting. Organoids were pelleted at 500 x g for 4 min at 4 °C, resuspended in TrypLE Express and dissociated for 15 min at 37 °C. Cells were counted and 10,000 cells were seeded per Matrigel dome on a 24-well plate.

### TSC derivations

Derivations of TSCs from human blastocysts^20^: Acidic Tyrode’s solution (T1788, Sigma) was used to remove the zona pellucida of human blastocysts as described above. The blastocysts devoid of zona pellucida were seeded on plates coated overnight with 5 µg/mL collagen IV (354233, Corning) in PBS and cultured in TSC medium for 4-5 days. The resulting TSC outgrowth was dissociated using TryplE for 15 min at 37 °C. The dissociated cells were plated on a new collagen IV coated dish and 0.25 µg/mL of iMatrix-511 (T303, Takara Bio) was added fresh to the media. After this step, we did not manage to recover TSC colonies.

Derivations of TSCs from TOs: TOs were dissociated into single cells as described above and seeded on 6-well plates coated overnight with 5 µg/mL collagen IV (354233, Corning) in PBS at 100,000 cells per well in TSC medium. iMatrix-511 (T303, Takara Bio) was added fresh to the media at a concentration of 0.25 µg/mL and media was changed every two to three days. Cells were cultured at 37 °C in 21% O_2_ and 5% CO_2_. When cultures reached ∼80% confluency, cells were dissociated with TrypLE for 15 min at 37 °C, pelleted at 500 x g for 4 min and passaged at a ratio of 1:8-1:16 depending on the confluency and growth of the cultures. After 1-2 passages in 2D with TSC medium, cultures were homogeneous with TSC morphology and could be maintained for over 40 passages.

TSC medium contains DMEM/F12 (21331020, Thermo Fisher Scientific), 1X GlutaMAX (35050061, Thermo Fisher Scientific), 1X penicillin-streptomycin (in house), 1X ITS-X (51500056, Thermo Fisher Scientific), 1% KnockOut Serum Replacement (10828028, Thermo Fisher Scientific), 0.15% BSA (A3311, Sigma-Aldrich), 800 µM valproic acid (P4543, Sigma-Aldrich), 200 µM L-ascorbic acid (A8960, Sigma-Aldrich), 25 ng/mL human EGF (Qk011, Qkine), 5 µM A83-01 (72022, STEMCELL Technologies), 2.5 µM Y27632 (Y-27632, STEMCELL Technologies), and 2 µM CHIR99021 (72054, STEMCELL Technologies).

### EVT differentiation

2D TSCs were differentiated to EVT as described previously^20^, with minor modifications. Briefly, 125,000 TSCs were seeded per well of a 6-well plate coated with 1 µg/mL collagen IV (354233, Corning) and cultured in 2D-EVT medium supplemented with 2% growth factor reduced Matrigel (356231, Corning). After three days, media was changed to EVT medium without NRG1 containing 0.5% growth factor-reduced Matrigel. At day six, media was replaced with EVT medium without NRG1 and Knockout Serum Replacement (KSR, 10828028, Thermo Fisher Scientific) containing 0.5% growth factor-reduced Matrigel. Cells were collected on day eight of the differentiation and analysed by flow cytometry or scRNA-seq.

2D-EVT medium contains DMEM/F12 (21331020, Thermo Fisher Scientific), 4% KnockOut Serum Replacement (10828028, Thermo Fisher Scientific), 1X GlutaMAX (35050061, Thermo Fisher Scientific), 1X penicillin-streptomycin (in house), 1X ITS-X (51500056, Thermo Fisher Scientific), 0.3% BSA (A3311, Sigma-Aldrich), 0.1 mM 2-mercaptoethanol (31350010, Thermo Fisher Scientific), 7.5 µM A83-01 (72022, STEMCELL Technologies), 2.5 µM Y27632 (Y-27632, STEMCELL Technologies) and 100 ng/mL NRG1 (100-03, Peprotech).

3D TOs were differentiated to EVT as described previously^13^, with minor modifications. After passaging, TOs were cultured in TOM for four to six days to allow recovery from single-cell dissociation before media was changed to 3D-EVT medium. Once cells started invading the Matrigel (ca. six days), the medium was changed to 3D-EVT medium without NRG1 and TOs were maintained for seven to ten days more before analysis by flow cytometry or scRNA-seq.

3D-EVT medium contains Advanced DMEM/F12 (12634010, Thermo Fisher Scientific), 4% KnockOut Serum Replacement (10828028, Thermo Fisher Scientific), 1X GlutaMAX (35050061, Thermo Fisher Scientific), 1X penicillin-streptomycin (in house), 1X ITS-X (51500056, Thermo Fisher Scientific), 0.3% BSA (A3311, Sigma-Aldrich), 0.1 mM 2-mercaptoethanol (31350010, Thermo Fisher Scientific), 7.5 µM A83-01 (72022, STEMCELL Technologies) and 100 ng/mL NRG1 (100-03, Peprotech).

### ExM derivation and maintenance

During TO derivation from entire embryos as well as derivations from polar halves and inner cells following immunosurgery, mesenchymal cells can sometimes be observed during the initial passages, usually alongside TOs. During routine passaging these cells are easily lost, as they do not form a compact pellet while TOs do. To maintain ExM cells, after 20 min Dispase treatment and centrifugation at 500 x g for 4 min at 4 °C, the supernatant was carefully removed with a pipette to ensure the mesenchymal cells were not lost. A loose cloud may be present above the pellet after centrifugation and care should be taken not to aspirate this, which usually meant leaving 100-200 µL liquid in the tube. The cells were resuspended in 700 µL dissociation solution containing DMEM/F12 (21331-020, Thermo Fisher Scientific), 1 mg/mL collagenase I (C2674, Sigma-Aldrich), 1 mg/mL collagenase IV (17104019, Gibco) and 20 U/mL DNase I (M0303S, New England Biolabs) and incubated for 15 min at 37 °C. The cells were washed with DMEM/F12 containing 0.1% BSA (A3311, Sigma-Aldrich), counted and 20,000 cells were seeded per 40 µL Matrigel dome on a 24- well plate in TOM.

For 2D culture, following single-cell dissociation of 3D ExM, cells were seeded on 6-well plates coated overnight with 5 µg/mL collagen IV (354233, Corning) in PBS at 100,000 cells per well in 2D-ExM medium. Media was changed every two to three days. When cultures reached ∼80% confluency, cells were dissociated with TrypLE for 15 min at 37 °C, pelleted at 500 x g for 4 min and passaged at a ratio of 1:2-1:8 depending on the confluency and growth of the cultures.

2D-ExM medium contains DMEM/F12, 1X GlutaMAX, 1X penicillin-streptomycin (in house), 1X ITS-X (51500056, Thermo Fisher Scientific), 0.3% BSA, 0.2% FBS (10270-106, Thermo Fisher Scientific), 0.1 mM 2-mercaptoethanol (31350010, Thermo Fisher Scientific), 1.5 µg/mL L-ascorbic acid (A8960, Sigma-Aldrich), 800 µM valproic acid (P4543, Sigma-Aldrich), 50 ng/mL human EGF (Qk011, Qkine), 5 µM Y27632 (Y-27632, STEMCELL Technologies), 2 µM CHIR99021 (72054, STEMCELL Technologies), 1 µM SB431542 (72232, STEMCELL Technologies), and 0.5 µM A83-01 (72022, STEMCELL Technologies).

### Phase-contrast microscopy

Phase-contrast images of live cells were taken on an EVOS FL (Invitrogen) and processed using Fiji 2.16.0 (https://fiji.sc/).

### Fluorescence activated cell sorting

To establish homogeneous ExM and TO cultures, early heterogeneous cultures were sorted using conjugated antibodies against E-CAD and ITGA6 (Table S6). After single-cell dissociation, cells were incubated in FACS buffer (PBS + 5% FBS (10270-106, Thermo Fisher Scientific)) and primary antibodies for one hour on ice. Next, cells were washed three times in FACS buffer, filtered using a 70 µm cell strainer and sorted into double-negative and double-positive populations on a FACSAria Fusion (BD Biosciences) flow cytometer. For recovery, 20,000 cells were seeded per 40 µL Matrigel dome on a 24-well plate in TOM.

### Flow cytometry

For HLA-G profiling of TSCs and TOs during EVT differentiation, single-cell suspensions were prepared in FACS buffer and incubated with a conjugated primary antibody against HLA-G (Table S6) for one hour on ice. Following incubation, cells were washed thrice in FACS buffer, filtered with a 70 µm cell strainer and analysed on a CytoFLEX flow cytometer (Beckman Coulter). Data analysis was carried out using FlowJo 10.10.0.

### Human ESC derivations and EB formation

The polar side of a human blastocyst was separated from the mural side as described above and plated on human foreskin fibroblast feeder cells in human ESC XenoFree culture medium at 37 °C in 5% O_2_ and 5% CO_2_. The XenoFree human ESC medium contains Knock-out DMEM (10829018, Thermo Fisher Scientific), 20% XenoFree KSR (12618012, Thermo Fisher Scientific), 1 mM GlutaMAX (35050061, Thermo Fisher Scientific), 0.1 mM 2- mercaptoethanol (M6250, Sigma), 10 mM non-essential amino acids (11140050, Thermo Fisher Scientific), and 4 ng/mL animal-free human recombinant basic FGF (GF003AF- 100UG, Merck). The outgrowth was manually split after five to seven days using a flame-polished borosilicate glass tool of approximately 100 µm size and expanded on feeders. After four passages on feeders, the human ESCs were cultured on Matrigel- (354277, Corning) coated plates with mTESR1 medium (85850, STEMCELL Technologies). Medium was changed daily and human ESCs were were mechanically passaged at 1:5-1:10 using L7 passaging solution (FP-5013, Lonza). Human ESCs were routinely cultured at 37 °C in 21% O_2_ and 5% CO_2_.

To generate EBs, colonies were mechanically cut as described above, and detached cell clumps were transferred to a 60 mm petri dish (353652, BD Falcon) containing 6 mL of AggreWell EB formation medium (05893, STEMCELL Technologies). The media was changed every other day, and after seven days of differentiation, EBs were collected for downstream analyses.

### Karyotyping by low coverage DNA sequencing

Genomic DNA was extracted from cell pellets using the Qiagen DNeasy Blood and Tissue kit (69506, Qiagen). Sequencing libraries were prepared with the NEBNext Ultra II kit (E6177L, New England Biolabs) and NEBNext Index Primers (E7335L, New England Biolabs) and were sequenced in house on a NextSeq 2000 (Illumina) with 0.3X coverage or higher. Reads were mapped to the hg38 reference genome with the Burrow-Wheeler Aligner^66^, sorted with Samtools^67^ and filtered for uniquely mapping reads using Sambamba^68^. Copy number analysis was carried out with the Control-FREEC package^69^ and results were visualised in R.

### Karyotyping by G-banding

G-banding was performed on 20 metaphase spreads of human ESCs at passage 10 by Cell Line Genetics (Madison, US). Metaphase spreads were imaged using a Leica GSL 120 CytoVision Scanner (Leica MicroSystems) with a 100x objective and a band count resolution of approximately 475.

### Immunofluorescence

TOs, TSCs and ExM cultures were fixed with 4% paraformaldehyde solution (28908, Thermo Fisher Scientific) at room temperature for 20 min. Samples were washed three times with PBST (PBS + 0.1% Tween-20 (P7949, Sigma Aldrich)) and permeabilised with PBS containing 0.3% Triton X-100 and 0.1 M glycine with for 30 min at room temperature. Next, samples were incubated in blocking buffer (PBS + 0.1% Tween-20 + 3% BSA) for 30 min at room temperature. Primary antibodies (Table S6) were diluted in blocking buffer were added overnight at 4 °C. Following three washes with PBST, samples were incubated with secondary antibody (Table S6) and DAPI (D1306, Thermo Fisher Scientific) at 1/500 dilution in blocking buffer overnight at 4 °C. Samples were washed thrice with PBST and imaged with an inverted SP8 confocal microscope (Leica Microsystems) with a 40x/1.1NA Water objective (Leica Microsystems). Images were processed using Fiji 2.16.0 (https://fiji.sc/).

Human embryos were stained in the same manner, with a few alterations. For fixations during TO derivation, embryos were removed from Matrigel domes using careful manual pipetting and 20 min Dispase treatment ahead of fixation with 4% paraformaldehyde. For VIM staining during TO derivation, embryos were permeabilised with PBS containing 0.5% Triton X-100 and 0.1 M glycine. Embryos were cleared using Refractive Index matching Solution (RIMS) buffer^70^. RIMS is composed of 2 g / mL of Histodenz (D2158, Sigma) dissolved in 0.02 M phosphate buffer (pH 7.4).

Human ESCs were fixed with a 4% paraformaldehyde (28908, Thermo Fisher Scientific) solution at room temperature for 15 minutes. Next, cells were permeabilised with a DPBS-based solution containing 0.1% Triton X-100 (T8787, Sigma Aldrich) and 0.1% sodium citrate (S4641, Sigma Aldrich) for 10 minutes at room temperature. This was followed by a blocking step using a buffer containing 10% Normal Donkey Serum (017000121, Jackson Immuno Research Labs) and 0.5% Triton X-100. Samples were incubated for 30 minutes at room temperature in blocking buffer. Primary antibody dilutions (Table S6) were prepared in blocking buffer and incubated overnight at 4 °C in a humid chamber. After three washes with DPBS, the fixed cells were incubated with fluorescent secondary antibody dilutions (Table S6) prepared in blocking buffer for 30 minutes at room temperature in a dark humid chamber. This was followed by three washes with DPBS. Cell nuclei were labelled with a 1/1000 dilution of Hoeschst 33342 (PI62249, Thermo Fisher Scientific). Samples were imaged with an Olympus IX71 microscope.

### Immunohistochemistry

TOs were fixed in 4% paraformaldehyde overnight at 4°C, paraffin embedded and sectioned at 6 µm intervals onto glass slides. TO slices underwent the following washes and stains; 3 x 5 minutes xylene, 3 minutes 100% ethanol, 3 minutes 95% ethanol, 3 minutes dH2O, 5min haematoxylin, 2 x 3 minutes dH2O, 20 seconds 0.1 HCl ethanol, 3 min dH2O, 2 x 2 minutes dH2O, 5 minutes eosin, 5 minutes dH2O, 2 x 10 seconds 100% ethanol, 3 minutes 100% ethanol, 5 minutes 100% ethanol, 3 x 5 minutes xylene. Slides were imaged with an AxioObserver inverted microscope (Zeiss) using the 20x Plan-Apochromat/0.80NA or 40x Plan-Apochromat oil/1.4NA objectives (Zeiss).

After removal of Matrigel with Dispase, ExM aggregates were fixed with 4% PFA (28908, Thermo Fisher Scientific) diluted in PBS for 20 minutes at room temperature. After three washes with PBS 0.1% Tween, the aggregates were embedded in Optimal Cutting Temperature Compound (OCT, 4583, Tissue-Tek). A cryostat was used to make 10-12 µm sections. These were stained using the immunofluorescence protocol outlined above. ProLong Gold Antifade Mountant with DNA stain (P36935, Thermo Fisher Scientific) was used as a mounting media and slides were sealed using CoverGrip Coverslip Sealant (23005, Biotium).

### Reverse transcription quantitative PCR of EBs

RNA from EBs was extracted using TRIzol Reagent (15596026, Thermo Fisher Scientific) and isolated using the Qiagen RNeasy Kit (74104, Qiagen) following the manufacturer’s instructions. The reverse transcription of total RNA to single-strand cDNA was achieved with the High-Capacity cDNA Reverse Transcription Kit (4368814, Applied Biosystems). Multi- lineage differentiation of EBs was analysed in triplicate by gene expression of molecular markers associated with the three somatic germ layers using reverse transcription quantitative PCR (1725272, SsoAdvanced Universal SYBR Green Supermix). ACTIN was used as the internal control. A list of the primers used is provided in Table S7.

### DNA methylation analysis

The bisulfite-pyrosequencing assay was designed using PyroMark Assay Design software (version 2.0) with primers provided in Table S7. The assay was validated using a standard curve calibrated against manufactured human genomic DNA 0% (80-8061-HGHM-5, EpigenDx) and 100% methylation controls (80-8062-HGUM-5, EpigenDx). Genomic DNA was first isolated from TOs using the DNeasy Blood and Tissue Kit (69504, Qiagen). Bisulfite conversion was then performed on 200-300 ng of genomic DNA from each sample using the EZ DNA Methylation-Gold Kit (D5006, Zymo Research). The bisulfite-converted DNA was amplified using HotStar Taq, and the PCR product was pyrosequenced on a PyroMark MD instrument (Qiagen). The methylation status of each CpG site was determined using the PyroMark Q-CpG software (Qiagen). The data from each CpG site (n = 6 per sample) was averaged and presented as a percent methylation for each sample.

### 10x genomics scRNA-seq

For scRNA-seq, TOs, TSCs, and ExM cells were dissociated into single cells as described above. For dome derivations, TOs and TSCs from embryo MS172 were used in the analysis, while the outgrowth dataset was generated from embryo MS104. TSCs were collected at 80% confluence. TOs were cultured in TOM or EVT differentiation conditions (six days in TOM, six days in 3D-EVT, ten days in 3D-EVT without NRG1) and collected 22 days after seeding. Cells were filtered using a 40 µm cell strainer and counted. Successful EVT differentiation was confirmed by flow cytometry for HLA-G. For ExM analysis, we used ExM derived from embryo MS615. 3D ExM was collected 13 days after plating, 2D ExM cells were collected after. Sequencing libraries were constructed using the Chromium Next GEM Single Cell 5’ Reagent Kit v2 (PN-1000263, 10X Genomics) according to the manufacturer’s instructions, with a target cell recovery of 10,000 cells per condition. Samples were sequenced on a NovaSeq 6000 or NextSeq 2000 at >50,000 reads per cell.

### scRNA-seq data processing

Analysis of trophoblast models:

Data repository integration: Droplet-based first-trimester scRNA-seq decidual and placental tissue data was obtained from the public Gene Expression Omnibus (GEO) repository (GEO174481 and GEO216244) and the ArrayExpress public repository (E-MTAB-6701). Raw BAM files from the GEO and ArrayExpress repositories were downloaded and pre- processed using Cell Ranger version 3.0 (10X Genomics). All samples underwent STAR read alignment as well as barcode and UMI counting against the hg19 reference genome.

Data pre-processing and quality control: To ensure only high-quality cells were used in downstream analyses, cells containing fewer than 500 detected genes and greater than 20% mitochondrial DNA content were removed. Individual samples were first scored based on expression levels of G2/M and S phase cell cycle gene markers. After this, individual samples were scaled, normalized, and contaminating cell doublets were removed using the scDblFinder package version 1.18.0^71^ and a doublet formation rate estimated from the number of captured cells. To minimize bias due to read dropouts, the “FindVariableFeatures” function was used with a “vst” selection method to prioritize highly variable genes in each sample for all downstream analyses. Following pre-processing, all samples were merged and subsequently integrated using cell pairwise correspondences between single cells across sequencing batches. During integration, the Pearson residuals obtained from the default regularized negative binomial model were re-computed and scaled to remove latent variation in percent mitochondrial DNA content as well as latent variation resulting from the difference between the G2/M and S phase cell cycle scores.

Trophoblast sub-setting, cell clustering, and identification: Trophoblasts were identified as fulfilling one of two criteria: First, trophoblasts were selected if they expressed combinations of *KRT7*, *EGFR*, *TEAD4*, *TP63*, *TFAP2A*, *TFAP2C*, *GATA2*, or *GATA3*, if they expressed EVT-enriched *HLA*-G, or if they expressed SCT-enriched *ERVFRD-1* transcripts at a level greater than zero. Trophoblasts further needed to express *VIM*, *WARS*, *PTPRC*, *DCN*, *CD34*, *CD14*, *CD86*, *CD163*, *NKG7*, *KLRD1*, *HLA*-*DPA1*, *HLA*-*DPB1*, *HLA-DRA*, *HLA-DRB1*, *HLA-DRB5*, and *HLA-DQA1* transcripts at a level equal to zero. Second, trophoblasts were selected if they originated from the villi TO, Okae TO, blastocyst TO, or the blastocyst TSC dataset, regardless of gene expression. After identification, trophoblasts were subset and re-clustered in Seurat at a resolution of 0.60 using 30 principal components. Cell identity was determined through observation of known trophoblast gene markers (*CGB*, *ERVFRD-1*, *HLA-G*, *ITGA5*, *EGFR*, *TP63*, *ELF5*, *BCAM*) as well as through observation of proliferative gene markers (*CCNA2* and *MKI67*) in each cluster and within each model. Cell state identity was further confirmed using cluster marker genes found via the “FindAllMarkers” function using a model-based analysis of single-cell transcriptomics (MAST) GLM-framework^72^ (v1.18.0), implemented in Seurat version 5.0.2^73^. Gene markers were selected as genes with a minimum log2 fold change value > 0.25 and an adjusted p-value < 0.05 within a specific cluster and are listed in Table S2. Trophoblast projections from all models were visualized together as an integrated data object using Uniform Manifold Approximation and Projections (UMAPs). This dataset is herein referred to as the integrated trophoblast dataset.

To independently compare each trophoblast model to the *in vivo* trophoblast reference data, primary placenta (chorionic villi) and decidua-derived trophoblasts were subset from the integrated dataset and re-clustered in Seurat using 50 principal components and are referred to as the *in vivo* dataset. Cell state identities were not re-calculated and are therefore retained from the integrated dataset. Trophoblast cells from *in vitro* models were similarly subset by model, and are herein referred to as villi TO, Okae TO, blastocyst TO and blastocyst TSC datasets respectively. Model-specific trophoblast projections were visualized using Uniform Manifold Approximation and Projections (UMAPs).

Pseudo-bulk Pearson’s correlation coefficient analysis: To specifically compare, without bias, the gene expression profile of CTB found in each trophoblast model to the uncultured CTB found *in vivo*, all CTB states were subset from the integrated trophoblast object, split by model, and CTB gene expression matrices were independently averaged by model (*in vivo* CTB, villi TO CTB, Okae TO CTB, blastocyst TO CTB, or 2D blastocyst TSC CTB) and log transformed to create model-specific “pseudo-bulk” CTB gene expression data. Pearson correlation coefficients of the relationship between these pseudo-bulk CTB expression profiles were calculated using the “cor” function from the stats base package available in R version 4.1.0. An identical approach was applied to compare, without bias, the gene expression profile of the cCTB, EVT, and SCTp states in each trophoblast model. Results from all comparisons were provided as tables (containing the raw data output) and as heatmaps generated using ggplot2 (version 3.5.0, https://ggplot2.tidyverse.org).

Pseudotime analysis: Monocle3 version 1.0.0^74^ was used to explore the differentiation of progenitor trophoblasts (i.e., CTB or TSC) into SCTp and EVT endpoints in the *in vivo* trophoblast reference and in each trophoblast model. In brief, expression matrices were input, clustered, and the principal graph was created using the “learn_graph” function. The “order_cells” function was then used to order cells in pseudotime without bias. The root node was computationally predicted using the “get_earliest_principal_node” function. Results for each model were visualized by model-specific UMAPs with the trajectory structure overlain.

Analysis of ExM cells:

Data pre-processing and quality control: Raw sequencing data were aligned using STARsolo (v2.7.10a_alpha_220818) against the GRCh38/hg38 human reference genome (Ensembl release 2020A). Gene expression matrices were obtained directly from STARsolo output using default parameters for the 10x 5′ v2 chemistry.

Putative doublets were identified using scrublet (version 0.2.3) applied individually to each sample. Cells were filtered to retain those expressing ≥2,000 genes and with <20% mitochondrial RNA content. Genes expressed in fewer than five cells were excluded. To control for batch imbalance, a maximum of 1,000 high-quality cells per sample were randomly retained for analysis.

Cell clustering and cell type annotation: Gene expression matrices were log-normalized using Scanpy (1.11.0). Highly variable genes (HVGs) were selected using the Seurat v3 algorithm (as implemented in Scanpy) with a batch-aware approach, selecting the top 4,000 HVGs across datasets using the dataset field as the batch key. This gene set was used in all downstream dimensionality reduction and correlation analyses.

Principal component analysis (PCA) was performed on log-normalized counts restricted to HVGs. The top 20 PCs, as determined by an elbow plot, were used to construct a k-nearest neighbor graph (k = 15). Cell clustering was performed using the Leiden algorithm.

Clusters were annotated manually based on curated marker gene lists and supported by published annotations from the original datasets. Marker genes used for annotation are provided in Table S5 and were informed by previous studies including Pham *et al*^7^. Additionally, cell type predictions for the invitro 2D and 3D ExM-like cells were conducted using the local version of Early Embryogenesis Prediction Tool (Version 2) as described https://zenodo.org/records/12189592.

Data integration with *in vivo* datasets: Human embryo *in vivo* sc-RNAseq datasets were obtained (Tyser *et al.*^50^: E-MTAB-9388, Xiang *et al.*^49^: GSE136447, Mole *et al.*^47^: E-MTAB- 8060, Ai *et al.*^48^: PRJCA017779) and processed as described above. Integration across all datasets was performed using harmonypy (version 0.0.10), with both dataset and sequencing method used as integration parameters. Harmony was run on the first 50 principal components with a theta parameter of 0 and a maximum of 30 iterations to ensure convergence.

Differential gene expression analysis: Differential expression between cell types was performed using the Wilcoxon rank-sum test as implemented in Scanpy’s rank_genes_groups function. Cell types were compared using the final manual annotations. Genes with log fold change > 0.25 and adjusted p-value < 0.05 were retained (Table S5).

Pseudo-bulk Pearson’s correlation coefficient analysis: To assess the similarity of transcriptional profiles across datasets, pearson’s correlations were calculated on the mean expression of HVGs. For each dataset and cell type combination, average gene expression values were computed. Pairwise Pearson’s correlation for the dataset-cell type combination were then calculated to generate a correlation heatmap.

### Statistical analyses

Statistical analyses were carried out using GraphPad Prism 10.4.2. Qualitative data was analysed using a two-sided Fisher’s exact test. Quantitative data was analysed with a one-way ANOVA test (multiple groups) or a two-tailed Student’s t-test (two groups). Human embryos that were dead at thawing or that did not recover 4 hours post-thawing were excluded from the study. Embryos that were accidentally damaged during dissections (e.g., polar/mural separation or immunosurgery) were also discarded.

## SUPPLEMENTARY FIGURE LEGENDS

Figure S1:Characterisation of blastocyst TSCs. **(A)** Schematic of the derivation protocol. **(B)** Phase-contrast images of Okae and blastocyst TSCs. Scale bars, 500 µm. **(C)** Immunofluorescence images of TSCs. Scale bars, 50 µm. **(D)** Phase- contrast images of EVT differentiation of blastocyst TSCs. Scale bars, 500 µm. **(E-F)** Flow cytometry of HLA-G levels of Okae and blastocyst TSCs in regenerative conditions and after EVT differentiation. Data is shown as mean ± SEM. n = 3 Okae TSC, n = 8 dome TSCs, n = 6 outgrowth TSCs, n = 3 Okae EVT, n = 10 dome EVT, n = 8 outgrowth EVT. One-way ANOVA, ns: non-significant, **p ≤ 0.01, ***p ≤ 0.001, ****p ≤ 0.0001. 6 independent experiments.

Figure S2: Single-cell RNA sequencing analysis of TOs. **(A)** UMAP plots separately clustered for each condition: *in vivo* villous trophoblast, villi TOs, Okae TOs, blastocyst TOs, and blastocyst TSCs. Cell type annotations are retained from Figure 2A. TO datasets contain both organoids in regenerative medium as well as after EVT differentiation, while blastocyst TSCs are only from regenerative conditions. **(B)** UMAP plots from A overlaid with Monocle3 pseudotime trajectories. **(C)** Dot plot of the percentage of cells expressing EVT genes and the average log expression in each group separated by cluster and trophoblast model. vEVT: villous EVT, iEVT: interstitial EVT, GC: placental bed giant cells, eEVT: endovascular EVT. **(D)** UMAP plot of TOs derived from blastocysts using the dome and outgrowth protocols as well as their corresponding undifferentiated TSCs. TO datasets contain both organoids in regenerative medium as well as after EVT differentiation. **(E)** Proportions of cells in each cluster in D by derivation protocol and growth condition. **(F)** Dot plot of the percentage of cells in the TSC, CTB1 and CTB2 clusters in D expressing trophoblast marker genes and their average log expression in dome and outgrowth TSCs.

Figure S3: Derivation of aneuploid TOs. (A, **B)** Copy number analysis of TOs at different passages. **(C, D)** Copy number analysis of TOs derived from PGT-A-tested blastocysts showing a non-concordance (C) and concordance (D) case. **(E)** Phase- contrast image of trisomy 13 TSCs derived from trisomy 13 TOs. Scale bar, 500 µm. **(F)** Immunofluorescence images of trisomy 13 TSCs. Scale bar, 50 µm. PGT-A: pre- implantation genetic testing for aneuploidy.

Figure S4: Derivation of stem cells from polar and mural halves of blastocysts. **(A)** Phase-contrast images of human ESC derivations. Scale bars, 100 µm. **(B)** Karyotype analysis of human ESCs derived in A. **(C)** Immunofluorescence images of human ESCs in A. Scale bars, 40 µm. **(D)** RT-qPCR analysis of embryoid bodies made from human ESCs derived in A. **(E, F)** Copy number analysis of TOs derived from the polar and mural halves of a human blastocyst.

Figure S5: Characterisation of blastocyst ExM cells. **(A)** Pregnancy test from ExM conditioned media. **(B)** Immunofluorescence image (maximum projection) of a section of 3D ExM. Scale bar, 100 µm. **(C)** Phase-contrast image of blastocyst ExM cultured in 2D. Scale bar, 500 µm. **(D)** Immunofluorescence images of ExM cultured in 2D. Scale bar, 100 µm. **(E, F)** Copy number analyses of a euploid (E) and trisomic (F) ExM line. **(G, H)** Copy number analyses of an ExM and a TO line derived from the same embryo.

Figure S6: Transcriptional profile of blastocyst ExM cells. **(A)** UMAP from Figure 5L with the cells for each embryo reference dataset highlighted and their respective original annotations^47–50^. **(B)** Dot plot of the percentage of cells expressing amnion, ExM, hypoblast and mesoderm marker genes and the average log expression in 2D and 3D blastocyst ExM (‘Schwarz 2D’ and ‘Schwarz 3D’), 2D ESC-derived ExM (‘Pham 2D’)^7^, ExM cells annotated by Ai *et al.* and the ExM and advanced mesoderm (AdvM) from Tyser et al^50^. **(C)** UMAP from Zhao et al^51^. integrating various human embryo scRNA-seq datasets into a reference. **(D)** UMAP with 3D blastocyst ExM projected onto the reference UMAP from C. **(E)** UMAP with 2D blastocyst ExM projected onto the reference UMAP from C. **(F)** Predicted cell types of 3D and 2D blastocyst ExM using the prediction tool from Zhao et al^51^.

## Notes

### Competing Interest Statement

The authors have declared no competing interest.

## REFERENCES

1. Niakan, K.K., Han, J., Pedersen, R.A., Simon, C., and Pera, R.A. (2012). Human pre-implantation embryo development. Development 139, 829–841. 10.1242/dev.060426.

2. Turco, M.Y., and Moffett, A. (2019). Development of the human placenta. Development 146. 10.1242/dev.163428.

3. Shahbazi, M.N., and Pasque, V. (2024). Early human development and stem cell-based human embryo models. Cell Stem Cell 31, 1398–1418. 10.1016/j.stem.2024.09.002.

4. Carter, A.M. (2012). Evolution of placental function in mammals: the molecular basis of gas and nutrient transfer, hormone secretion, and immune responses. Physiol Rev 92, 1543–1576. 10.1152/physrev.00040.2011.

5. Hemberger, M., Hanna, C.W., and Dean, W. (2020). Mechanisms of early placental development in mouse and humans. Nat Rev Genet 21, 27–43. 10.1038/s41576-019-0169-4.

6. Pollheimer, J., Vondra, S., Baltayeva, J., Beristain, A.G., and Knofler, M. (2018). Regulation of Placental Extravillous Trophoblasts by the Maternal Uterine Environment. Front Immunol 9, 2597. 10.3389/fimmu.2018.02597.

7. Pham, T.X.A., Panda, A., Kagawa, H., To, S.K., Ertekin, C., Georgolopoulos, G., van Knippenberg, S., Allsop, R.N., Bruneau, A., Chui, J.S., et al. (2022). Modeling human extraembryonic mesoderm cells using naive pluripotent stem cells. Cell Stem Cell 29, 1346–1365 e1310. 10.1016/j.stem.2022.08.001.

8. Nehme, E., Panda, A., Migeotte, I., and Pasque, V. (2025). Extra-embryonic mesoderm during development and in in vitro models. Development 152. 10.1242/dev.204624.

9. Hoffman, M.K. (2023). The great obstetrical syndromes and the placenta. BJOG 130 *Suppl 3*, 8–15. 10.1111/1471-0528.17613.

10. Siriwardena, D., Munger, C., Penfold, C., Kohler, T.N., Weberling, A., Linneberg-Agerholm, M., Slatery, E., Ellermann, A.L., Bergmann, S., Clark, S.J., et al. (2024). Marmoset and human trophoblast stem cells differ in signaling requirements and recapitulate divergent modes of trophoblast invasion. Cell Stem Cell 31, 1427–1446 e1428. 10.1016/j.stem.2024.09.004.

11. Roberts, R.M., Green, J.A., and Schulz, L.C. (2016). The evolution of the placenta. Reproduction 152, R179–189. 10.1530/REP-16-0325.

12. Haider, S., and Beristain, A.G. (2023). Human organoid systems in modeling reproductive tissue development, function, and disease. Hum Reprod 38, 1449–1463. 10.1093/humrep/dead085.

13. Turco, M.Y., Gardner, L., Kay, R.G., Hamilton, R.S., Prater, M., Hollinshead, M.S., McWhinnie, A., Esposito, L., Fernando, R., Skelton, H., et al. (2018). Trophoblast organoids as a model for maternal-fetal interactions during human placentation. Nature 564, 263–267. 10.1038/s41586-018-0753-3.

14. Haider, S., Meinhardt, G., Saleh, L., Kunihs, V., Gamperl, M., Kaindl, U., Ellinger, A., Burkard, T.R., Fiala, C., Pollheimer, J., et al. (2018). Self-Renewing Trophoblast Organoids Recapitulate the Developmental Program of the Early Human Placenta. Stem Cell Reports 11, 537–551. 10.1016/j.stemcr.2018.07.004.

15. van Rijn, B.v.O., D.; van Koetsveld, N.; Knapen, M.; Gribnau, J.; Schäffers, O. (2024). Generation of trophoblast organoids from chorionic villus sampling. Organoids 3, 54–66. 10.3390/organoids3010005.

16. Castel, G., Meistermann, D., Bretin, B., Firmin, J., Blin, J., Loubersac, S., Bruneau, A., Chevolleau, S., Kilens, S., Chariau, C., et al. (2020). Induction of Human Trophoblast Stem Cells from Somatic Cells and Pluripotent Stem Cells. Cell Rep 33, 108419. 10.1016/j.celrep.2020.108419.

17. Karvas, R.M., Khan, S.A., Verma, S., Yin, Y., Kulkarni, D., Dong, C., Park, K.M., Chew, B., Sane, E., Fischer, L.A., et al. (2022). Stem-cell-derived trophoblast organoids model human placental development and susceptibility to emerging pathogens. Cell Stem Cell 29, 810–825 e818. 10.1016/j.stem.2022.04.004.

18. Cameron, S.T. (2025). Early medical abortion. Best Pract Res Clin Obstet Gynaecol 99, 102588. 10.1016/j.bpobgyn.2025.102588.

19. Thomson, J.A., Itskovitz-Eldor, J., Shapiro, S.S., Waknitz, M.A., Swiergiel, J.J., Marshall, V.S., and Jones, J.M. (1998). Embryonic stem cell lines derived from human blastocysts. Science 282, 1145–1147.

20. Okae, H., Toh, H., Sato, T., Hiura, H., Takahashi, S., Shirane, K., Kabayama, Y., Suyama, M., Sasaki, H., and Arima, T. (2018). Derivation of Human Trophoblast Stem Cells. Cell Stem Cell 22, 50–63 e56. 10.1016/j.stem.2017.11.004.

21. Sheridan, M.A., Zhao, X., Fernando, R.C., Gardner, L., Perez-Garcia, V., Li, Q., Marsh, S.G.E., Hamilton, R., Moffett, A., and Turco, M.Y. (2021). Characterization of primary models of human trophoblast. Development 148. 10.1242/dev.199749.

22. Shannon, M.J., McNeill, G.L., Koksal, B., Baltayeva, J., Wachter, J., Castellana, B., Penaherrera, M.S., Robinson, W.P., Leung, P.C.K., and Beristain, A.G. (2024). Single-cell assessment of primary and stem cell-derived human trophoblast organoids as placenta-modeling platforms. Dev Cell 59, 776–792 e711. 10.1016/j.devcel.2024.01.023.

23. Wang, D., Cearlock, A., Lane, K., Xu, C., Jan, I., McCartney, S., Glass, I., McCoy, R., and Yang, M. (2025). Chromosomal instability in human trophoblast stem cells and placentas. Nat Commun 16, 3918. 10.1038/s41467-025-59245-9.

24. Starostik, M.R., Sosina, O.A., and McCoy, R.C. (2020). Single-cell analysis of human embryos reveals diverse patterns of aneuploidy and mosaicism. Genome Res. 10.1101/gr.262774.120.

25. Yang, M., Rito, T., Metzger, J., Naftaly, J., Soman, R., Hu, J., Albertini, D.F., Barad, D.H., Brivanlou, A.H., and Gleicher, N. (2021). Depletion of aneuploid cells in human embryos and gastruloids. Nat Cell Biol 23, 314–321. 10.1038/s41556-021-00660-7.

26. Kalousek, D.K., and Dill, F.J. (1983). Chromosomal mosaicism confined to the placenta in human conceptions. Science 221, 665–667. 10.1126/science.6867735.

27. Kalousek, D.K., Howard-Peebles, P.N., Olson, S.B., Barrett, I.J., Dorfmann, A., Black, S.H., Schulman, J.D., and Wilson, R.D. (1991). Confirmation of CVS mosaicism in term placentae and high frequency of intrauterine growth retardation association with confined placental mosaicism. Prenat Diagn 11, 743–750. 10.1002/pd.1970111002.

28. Amor, D.J., Neo, W.T., Waters, E., Heussler, H., Pertile, M., and Halliday, J. (2006). Health and developmental outcome of children following prenatal diagnosis of confined placental mosaicism. Prenat Diagn 26, 443–448. 10.1002/pd.1433.

29. Eggenhuizen, G.M., Go, A., Koster, M.P.H., Baart, E.B., and Galjaard, R.J. (2021). Confined placental mosaicism and the association with pregnancy outcome and fetal growth: a review of the literature. Hum Reprod Update 27, 885–903. 10.1093/humupd/dmab009.

30. Fernandez Gallardo, E.S., A.; Chappell, J.; Demeulemeester, J.; Herrmann, J.C.; Vermotte, R.; Kerremans, A.; Van der Haegen, M.; Van Herck, J.; Vanuytven, S.; Vandereyken, K.; Macaulay, I.C.; Vermeesch, J.R.; Peeraer, K.; Debrock, S.; Pasque, V.; Voet, T. (2023). A multi-omics genome-and-transcriptome single-cell atlas of human preimplantation embryogenesis reveals the cellular and molecular impact of chromosome instability. bioRxiv.

31. Hemberger, M., Udayashankar, R., Tesar, P., Moore, H., and Burton, G.J. (2010). ELF5-enforced transcriptional networks define an epigenetically regulated trophoblast stem cell compartment in the human placenta. Hum Mol Genet 19, 2456–2467. 10.1093/hmg/ddq128.

32. Lee, C.Q., Gardner, L., Turco, M., Zhao, N., Murray, M.J., Coleman, N., Rossant, J., Hemberger, M., and Moffett, A. (2016). What Is Trophoblast? A Combination of Criteria Define Human First-Trimester Trophoblast. Stem Cell Reports 6, 257–272. 10.1016/j.stemcr.2016.01.006.

33. (!!! INVALID CITATION !!! 20).

34. Shannon, M.J., Baltayeva, J., Castellana, B., Wachter, J., McNeill, G.L., Yoon, J.S., Treissman, J., Le, H.T., Lavoie, P.M., and Beristain, A.G. (2022). Cell trajectory modeling identifies a primitive trophoblast state defined by BCAM enrichment. Development 149. 10.1242/dev.199840.

35. van Echten-Arends, J., Mastenbroek, S., Sikkema-Raddatz, B., Korevaar, J.C., Heineman, M.J., van der Veen, F., and Repping, S. (2011). Chromosomal mosaicism in human preimplantation embryos: a systematic review. Hum Reprod Update 17, 620–627. 10.1093/humupd/dmr014.

36. Coorens, T.H.H., Oliver, T.R.W., Sanghvi, R., Sovio, U., Cook, E., Vento-Tormo, R., Haniffa, M., Young, M.D., Rahbari, R., Sebire, N., et al. (2021). Inherent mosaicism and extensive mutation of human placentas. Nature 592, 80–85. 10.1038/s41586-021-03345-1.

37. Shahbazi, M.N., Wang, T., Tao, X., Weatherbee, B.A.T., Sun, L., Zhan, Y., Keller, L., Smith, G.D., Pellicer, A., Scott, R.T., Jr., et al. (2020). Developmental potential of aneuploid human embryos cultured beyond implantation. Nat Commun 11, 3987. 10.1038/s41467-020-17764-7.

38. Yong, P.J., Barrett, I.J., Kalousek, D.K., and Robinson, W.P. (2003). Clinical aspects, prenatal diagnosis, and pathogenesis of trisomy 16 mosaicism. J Med Genet 40, 175–182. 10.1136/jmg.40.3.175.

39. Grati, F.R., Ferreira, J., Benn, P., Izzi, C., Verdi, F., Vercellotti, E., Dalpiaz, C., D’Ajello, P., Filippi, E., Volpe, N., et al. (2020). Outcomes in pregnancies with a confined placental mosaicism and implications for prenatal screening using cell-free DNA. Genet Med 22, 309–316. 10.1038/s41436-019-0630-y.

40. Malvestiti, F., Agrati, C., Grimi, B., Pompilii, E., Izzi, C., Martinoni, L., Gaetani, E., Liuti, M.R., Trotta, A., Maggi, F., et al. (2015). Interpreting mosaicism in chorionic villi: results of a monocentric series of 1001 mosaics in chorionic villi with follow-up amniocentesis. Prenat Diagn 35, 1117–1127. 10.1002/pd.4656.

41. Biancotti, J.C., Narwani, K., Buehler, N., Mandefro, B., Golan-Lev, T., Yanuka, O., Clark, A., Hill, D., Benvenisty, N., and Lavon, N. (2010). Human embryonic stem cells as models for aneuploid chromosomal syndromes. Stem Cells 28, 1530–1540. 10.1002/stem.483.

42. McCoy, R.C., Demko, Z.P., Ryan, A., Banjevic, M., Hill, M., Sigurjonsson, S., Rabinowitz, M., and Petrov, D.A. (2015). Evidence of Selection against Complex Mitotic-Origin Aneuploidy during Preimplantation Development. PLoS Genet 11, e1005601. 10.1371/journal.pgen.1005601.

43. Rodriguez-Purata, J., Lee, J., Whitehouse, M., Moschini, R.M., Knopman, J., Duke, M., Sandler, B., and Copperman, A. (2015). Embryo selection versus natural selection: how do outcomes of comprehensive chromosome screening of blastocysts compare with the analysis of products of conception from early pregnancy loss (dilation and curettage) among an assisted reproductive technology population? Fertil Steril 104, 1460–1466 e1461-1412. 10.1016/j.fertnstert.2015.08.007.

44. Meistermann, D., Bruneau, A., Loubersac, S., Reignier, A., Firmin, J., Francois-Campion, V., Kilens, S., Lelievre, Y., Lammers, J., Feyeux, M., et al. (2021). Integrated pseudotime analysis of human pre-implantation embryo single-cell transcriptomes reveals the dynamics of lineage specification. Cell Stem Cell 28, 1625–1640 e1626. 10.1016/j.stem.2021.04.027.

45. Liu, D., Chen, Y., Ren, Y., Yuan, P., Wang, N., Liu, Q., Yang, C., Yan, Z., Yang, M., Wang, J., et al. (2022). Primary specification of blastocyst trophectoderm by scRNA-seq: New insights into embryo implantation. Sci Adv 8, eabj3725. 10.1126/sciadv.abj3725.

46. James, J.L., Carter, A.M., and Chamley, L.W. (2012). Human placentation from nidation to 5 weeks of gestation. Part I: What do we know about formative placental development following implantation? Placenta 33, 327–334. 10.1016/j.placenta.2012.01.020.

47. Mole, M.A., Coorens, T.H.H., Shahbazi, M.N., Weberling, A., Weatherbee, B.A.T., Gantner, C.W., Sancho-Serra, C., Richardson, L., Drinkwater, A., Syed, N., et al. (2021). A single cell characterisation of human embryogenesis identifies pluripotency transitions and putative anterior hypoblast centre. Nat Commun 12, 3679. 10.1038/s41467-021-23758-w.

48. Ai, Z., Niu, B., Yin, Y., Xiang, L., Shi, G., Duan, K., Wang, S., Hu, Y., Zhang, C., Zhang, C., et al. (2023). Dissecting peri-implantation development using cultured human embryos and embryo-like assembloids. Cell Res 33, 661–678. 10.1038/s41422-023-00846-8.

49. Xiang, L., Yin, Y., Zheng, Y., Ma, Y., Li, Y., Zhao, Z., Guo, J., Ai, Z., Niu, Y., Duan, K., et al. (2019). A developmental landscape of 3D-cultured human pre-gastrulation embryos. Nature. 10.1038/s41586-019-1875-y.

50. Tyser, R.C.V., Mahammadov, E., Nakanoh, S., Vallier, L., Scialdone, A., and Srinivas, S. (2021). Single-cell transcriptomic characterization of a gastrulating human embryo. Nature 600, 285–289. 10.1038/s41586-021-04158-y.

51. Zhao, C., Plaza Reyes, A., Schell, J.P., Weltner, J., Ortega, N.M., Zheng, Y., Bjorklund, A.K., Baque-Vidal, L., Sokka, J., Trokovic, R., et al. (2025). A comprehensive human embryo reference tool using single-cell RNA-sequencing data. Nat Methods 22, 193–206. 10.1038/s41592-024-02493-2.

52. Simon, C.S.M., A.; Woods, L.; Staneva, D.; Huang, Q.; Linneberg-Agerholm, M.; Faulkner, A.; Papathanasiou, A.; Elder, K.; Snell, P.; Christie, L.; Garcia, P.; Shaikly, V.; Taranissi, M.; Choudhary, M.; Herbert, M.; Brickman, J.M.; Niakan, K.K. (2024). Suppression of ERK signalling promotes pluripotent epiblast in the human blastocyst. bioRxiv.

53. Niakan, K.K., and Eggan, K. (2013). Analysis of human embryos from zygote to blastocyst reveals distinct gene expression patterns relative to the mouse. Dev Biol 375, 54–64. 10.1016/j.ydbio.2012.12.008.

54. Guo, G., Stirparo, G.G., Strawbridge, S.E., Spindlow, D., Yang, J., Clarke, J., Dattani, A., Yanagida, A., Li, M.A., Myers, S., et al. (2021). Human naive epiblast cells possess unrestricted lineage potential. Cell Stem Cell 28, 1040–1056 e1046. 10.1016/j.stem.2021.02.025.

55. Corujo-Simon, E., Bates, L.E., Yanagida, A., Jones, K., Clark, S., von Meyenn, F., Reik, W., and Nichols, J. (2024). Human trophectoderm becomes multi-layered by internalization at the polar region. Dev Cell 59, 2497–2505 e2494. 10.1016/j.devcel.2024.05.028.

56. Linneberg-Agerholm, M., Sell, A.C., Redo-Riveiro, A., Perera, M., Proks, M., Knudsen, T.E., Barral, A., Manzanares, M., and Brickman, J.M. (2024). The primitive endoderm supports lineage plasticity to enable regulative development. Cell 187, 4010–4029 e4016. 10.1016/j.cell.2024.05.051.

57. Enders, A.C., and King, B.F. (1988). Formation and differentiation of extraembryonic mesoderm in the rhesus monkey. Am J Anat 181, 327–340. 10.1002/aja.1001810402.

58. Luckett, W.P. (1978). Origin and differentiation of the yolk sac and extraembryonic mesoderm in presomite human and rhesus monkey embryos. Am J Anat 152, 59–97. 10.1002/aja.1001520106.

59. Siegel, J.J., and Amon, A. (2012). New insights into the troubles of aneuploidy. Annu Rev Cell Dev Biol 28, 189–214. 10.1146/annurev-cellbio-101011-155807.

60. Chavli, E.A., Klaasen, S.J., Van Opstal, D., Laven, J.S., Kops, G.J., and Baart, E.B. (2024). Single-cell DNA sequencing reveals a high incidence of chromosomal abnormalities in human blastocysts. J Clin Invest 134. 10.1172/JCI174483.

61. Benn, P. (1998). Trisomy 16 and trisomy 16 Mosaicism: a review. Am J Med Genet 79, 121–133.

62. Yang, L., Liang, P., Yang, H., and Coyne, C.B. (2024). Trophoblast organoids with physiological polarity model placental structure and function. J Cell Sci 137. 10.1242/jcs.261528.

63. Hori, T., Okae, H., Shibata, S., Kobayashi, N., Kobayashi, E.H., Oike, A., Sekiya, A., Arima, T., and Kaji, H. (2024). Trophoblast stem cell-based organoid models of the human placental barrier. Nat Commun 15, 962. 10.1038/s41467-024-45279-y.

64. Zhou, J., Sheridan, M.A., Tian, Y., Dahlgren, K.J., Messler, M., Peng, T., Zhao, A., Ezashi, T., Schulz, L.C., Ulery, B.D., et al. (2025). Development of apical out trophoblast stem cell derived organoids to model early human pregnancy. iScience 28, 112099. 10.1016/j.isci.2025.112099.

65. Strawbridge, S.E., Clarke, J., Guo, G., and Nichols, J. (2022). Deriving Human Naive Embryonic Stem Cell Lines from Donated Supernumerary Embryos Using Physical Distancing and Signal Inhibition. Methods Mol Biol 2416, 1–12. 10.1007/978-1-0716-1908-7_1.

66. Li, H. (2013). Aligning sequence reads, clone sequences and assembly contigs with BWA-MEM. arXiv. 10.48550/arXiv.1303.3997.

67. Danecek, P., Bonfield, J.K., Liddle, J., Marshall, J., Ohan, V., Pollard, M.O., Whitwham, A., Keane, T., McCarthy, S.A., Davies, R.M., and Li, H. (2021). Twelve years of SAMtools and BCFtools. Gigascience 10. 10.1093/gigascience/giab008.

68. Tarasov, A., Vilella, A.J., Cuppen, E., Nijman, I.J., and Prins, P. (2015). Sambamba: fast processing of NGS alignment formats. Bioinformatics 31, 2032–2034. 10.1093/bioinformatics/btv098.

69. Boeva, V., Popova, T., Bleakley, K., Chiche, P., Cappo, J., Schleiermacher, G., Janoueix-Lerosey, I., Delattre, O., and Barillot, E. (2012). Control-FREEC: a tool for assessing copy number and allelic content using next-generation sequencing data. Bioinformatics 28, 423–425. 10.1093/bioinformatics/btr670.

70. Yang, B., Treweek, J.B., Kulkarni, R.P., Deverman, B.E., Chen, C.K., Lubeck, E., Shah, S., Cai, L., and Gradinaru, V. (2014). Single-cell phenotyping within transparent intact tissue through whole-body clearing. Cell 158, 945–958. 10.1016/j.cell.2014.07.017.

71. Germain, P.L., Lun, A., Garcia Meixide, C., Macnair, W., and Robinson, M.D. (2021). Doublet identification in single-cell sequencing data using scDblFinder. F1000Res 10, 979. 10.12688/f1000research.73600.2.

72. Finak, G., McDavid, A., Yajima, M., Deng, J., Gersuk, V., Shalek, A.K., Slichter, C.K., Miller, H.W., McElrath, M.J., Prlic, M., et al. (2015). MAST: a flexible statistical framework for assessing transcriptional changes and characterizing heterogeneity in single-cell RNA sequencing data. Genome Biol 16, 278. 10.1186/s13059-015-0844-5.

73. Hao, Y., Stuart, T., Kowalski, M.H., Choudhary, S., Hoffman, P., Hartman, A., Srivastava, A., Molla, G., Madad, S., Fernandez-Granda, C., and Satija, R. (2024). Dictionary learning for integrative, multimodal and scalable single-cell analysis. Nat Biotechnol 42, 293–304. 10.1038/s41587-023-01767-y.

74. Cao, J., Spielmann, M., Qiu, X., Huang, X., Ibrahim, D.M., Hill, A.J., Zhang, F., Mundlos, S., Christiansen, L., Steemers, F.J., et al. (2019). The single-cell transcriptional landscape of mammalian organogenesis. Nature 566, 496–502. 10.1038/s41586-019-0969-x.

