## Supplementary figures and images for "A blastocyst-derived *in vitro* model of the *human* chorion"

### Supplementary Figures

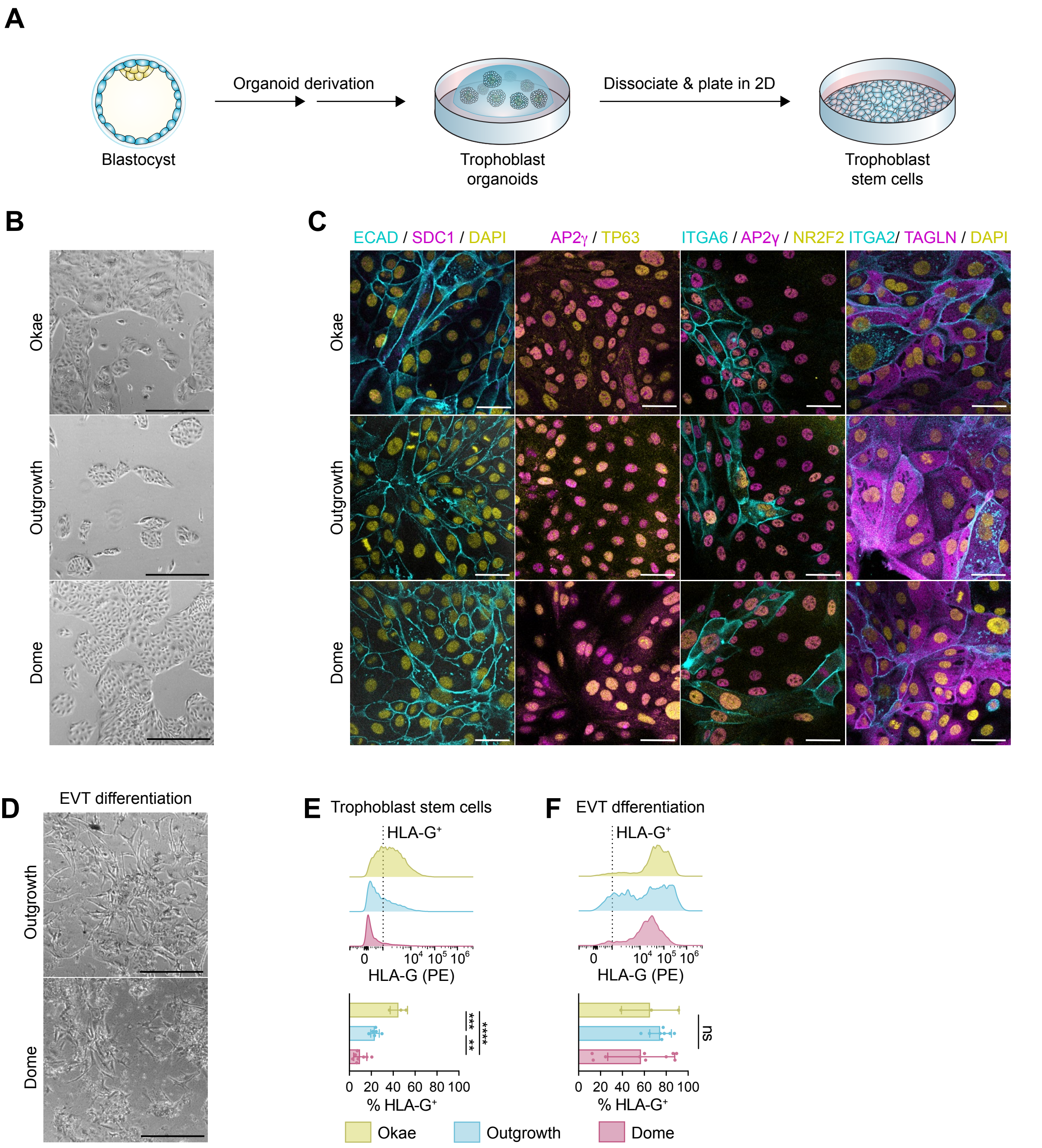

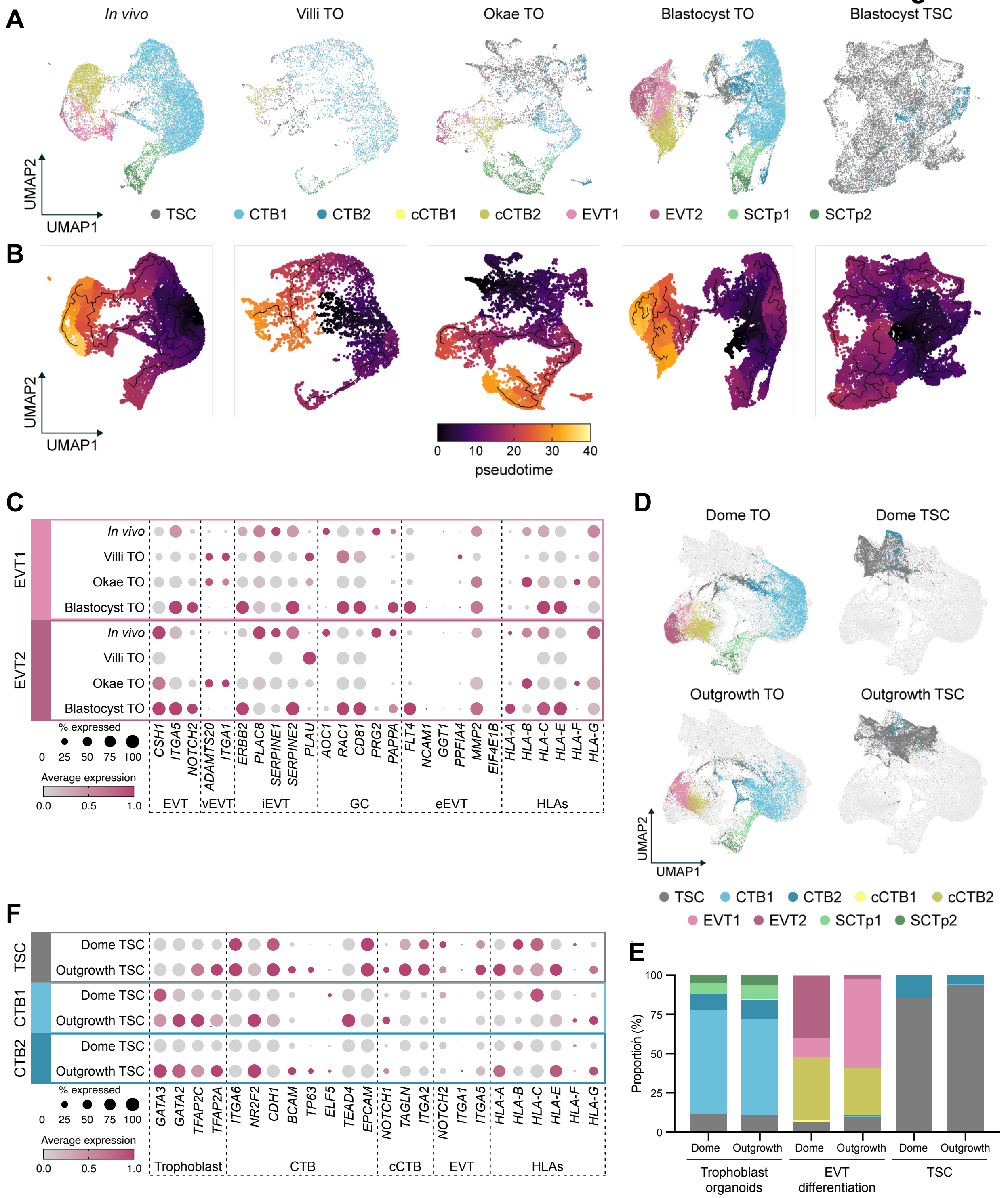

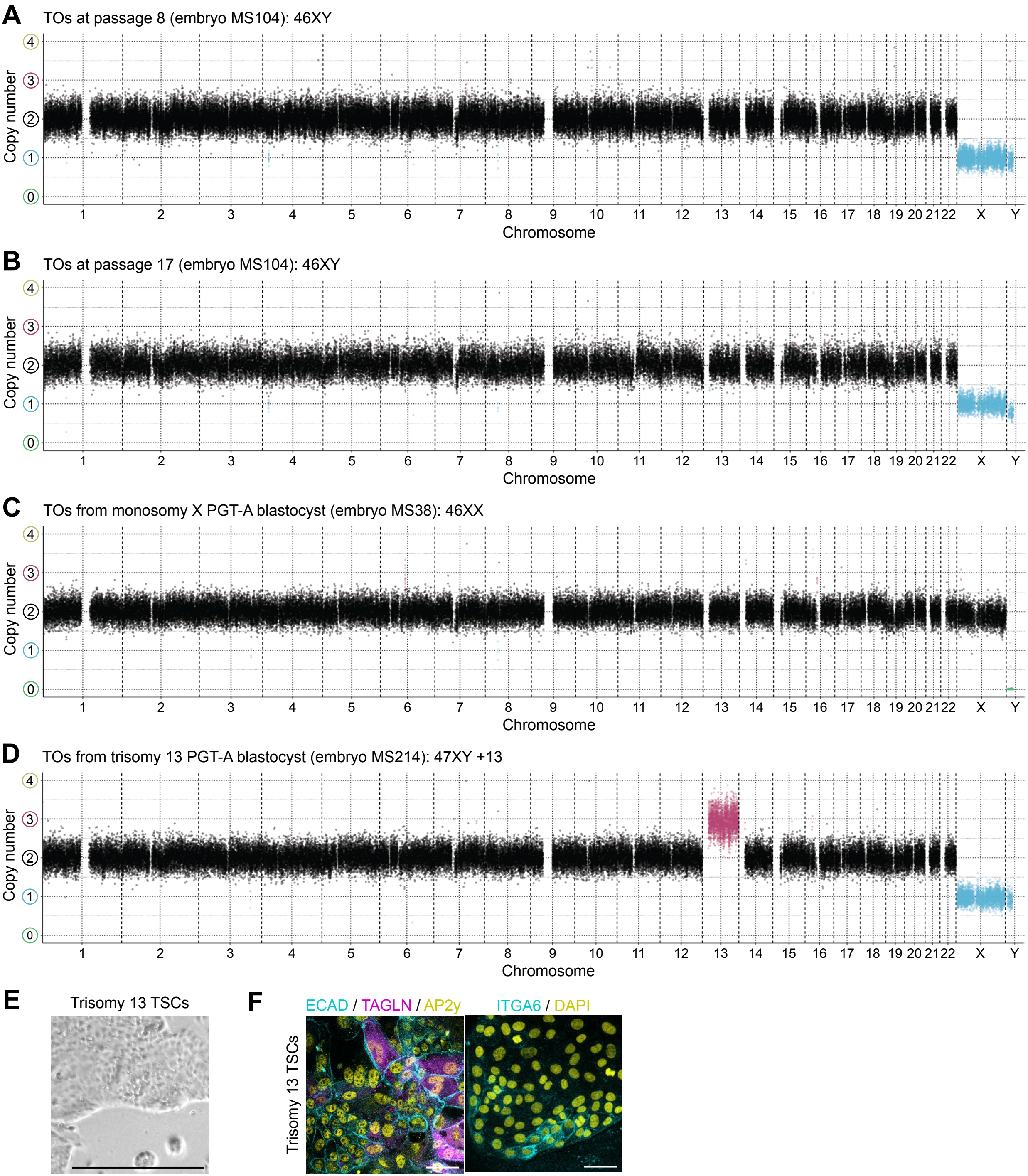

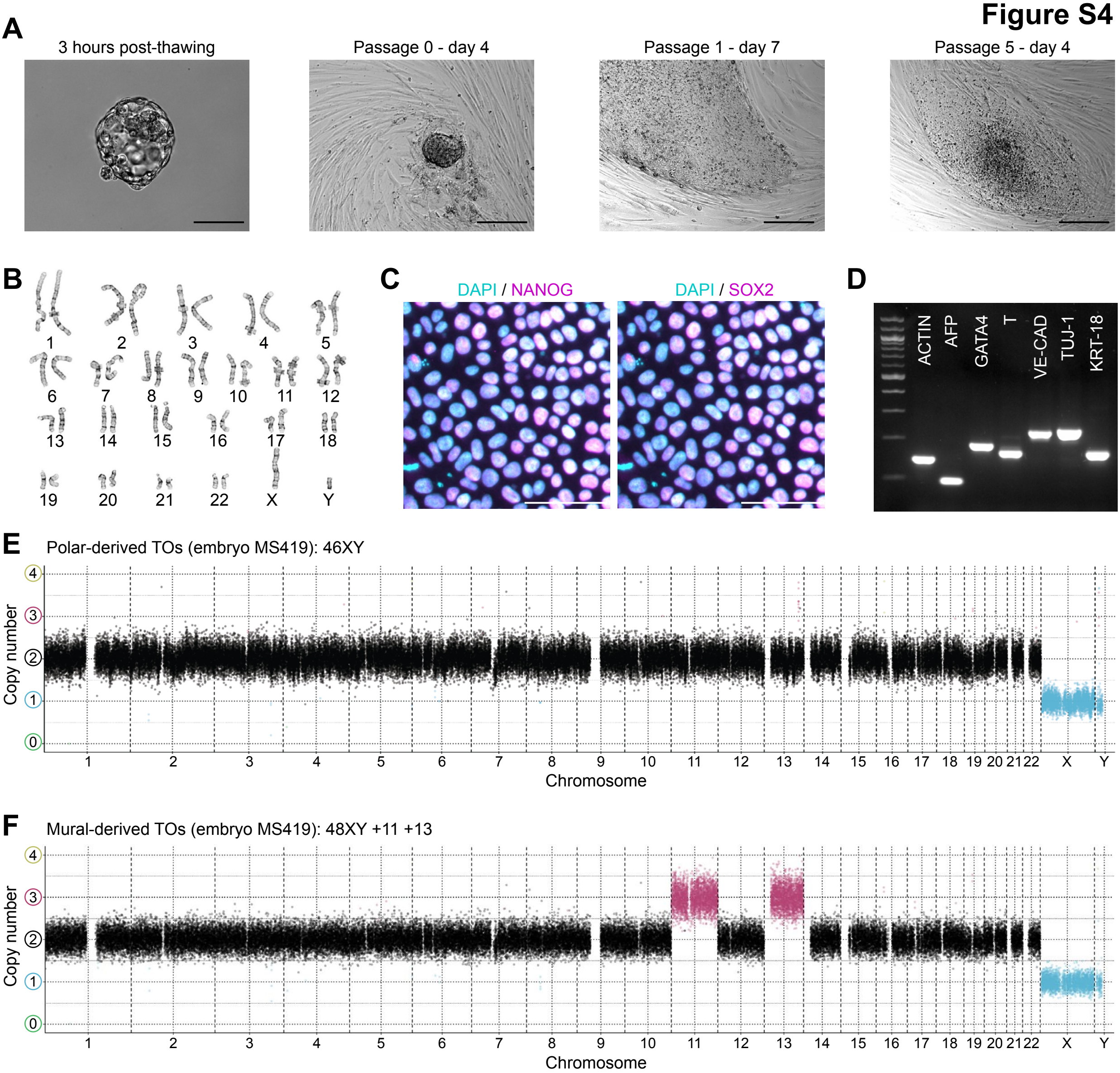

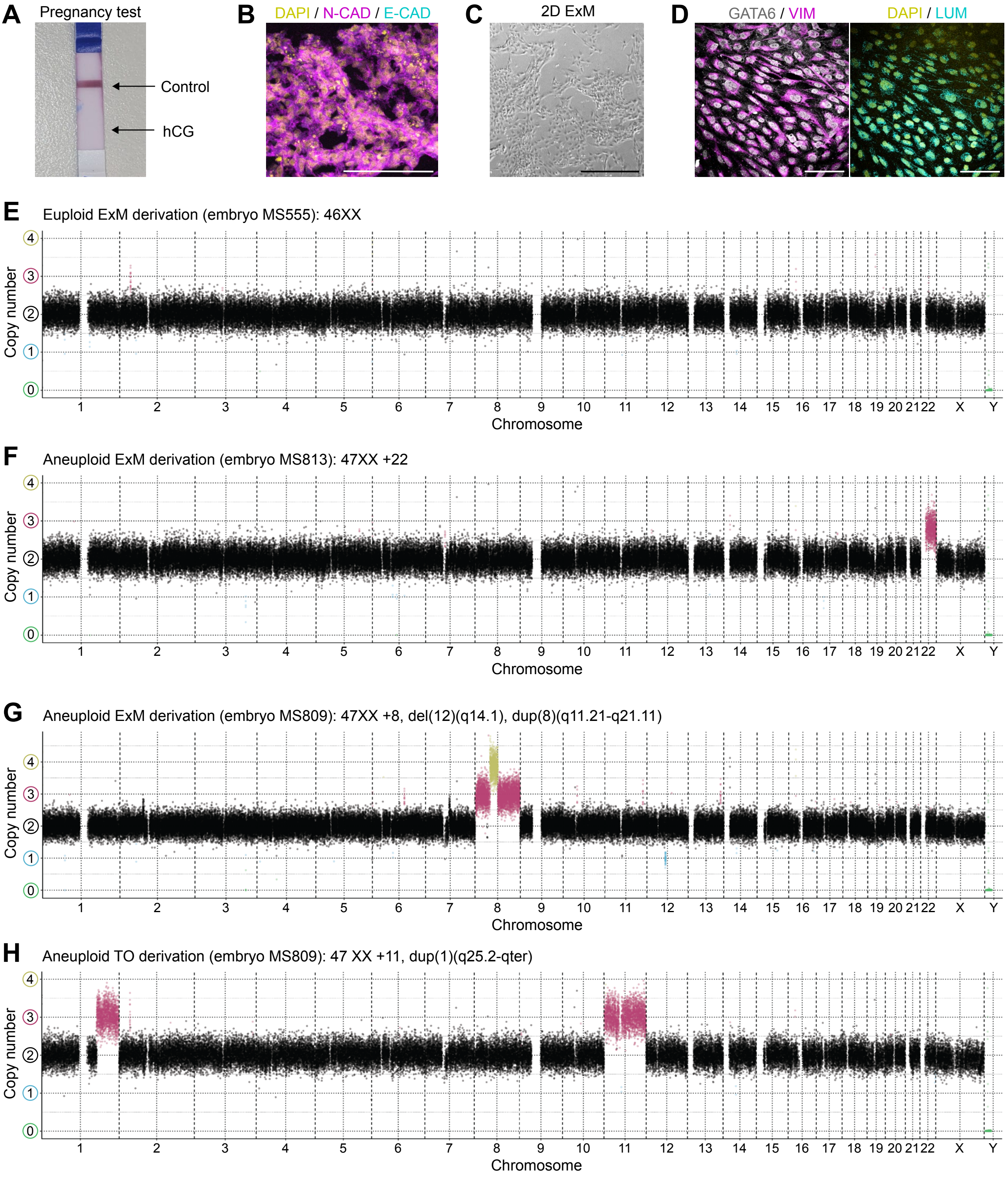

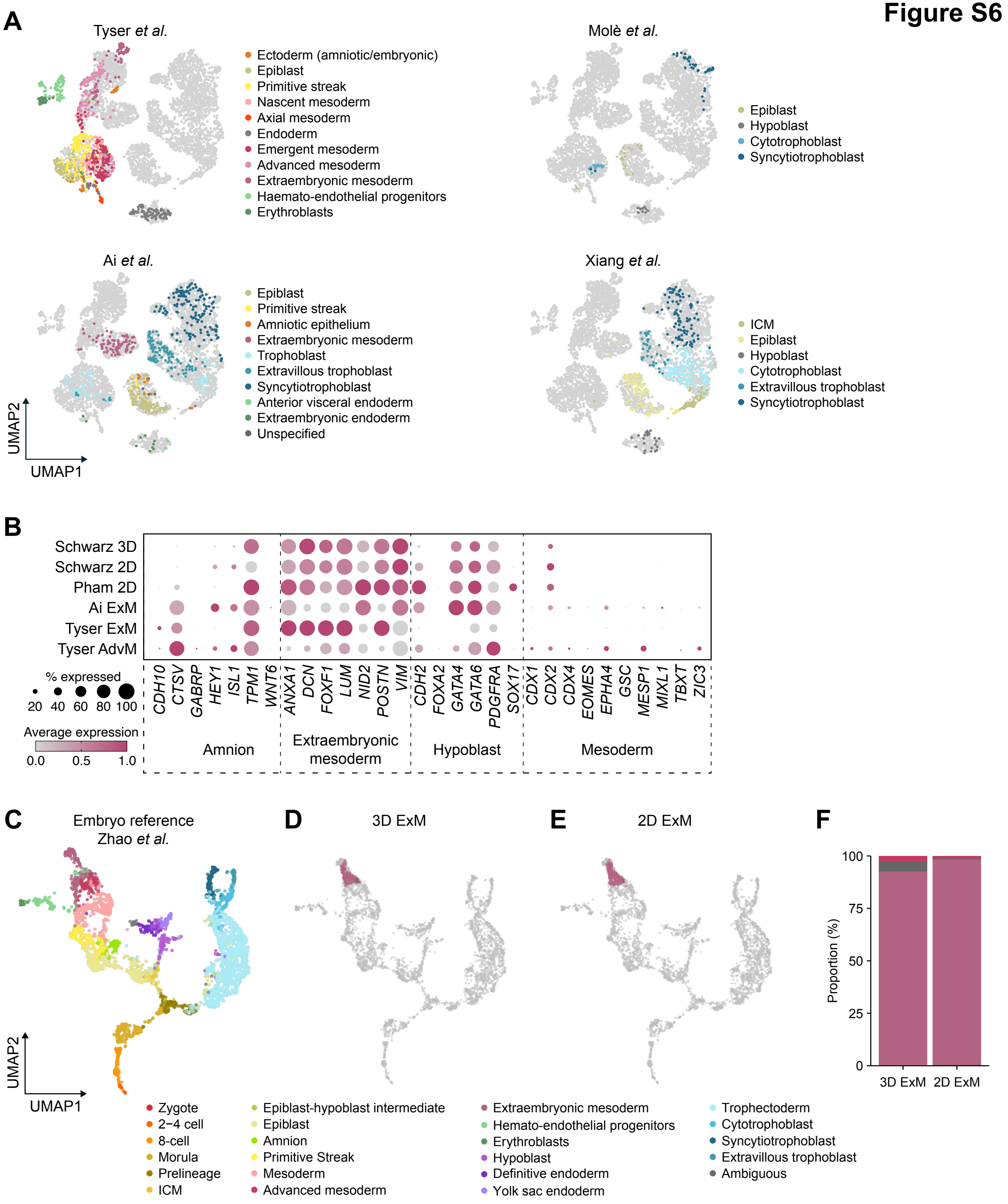
